## Supplementary Materials for "Whole-genome phylogenomics and synteny resolve a single origin of body-plan asymmetry in flatfishes"

### Outline

- Supplementary Tables
  - Supplementary Table S1. Specimen, voucher, sequencing-library, and accession metadata for the three newly assembled genomes.
  - Supplementary Table S2. Genome assembly statistics for representative assemblies of the three newly assembled genomes.
  - Supplementary Table S3. Assembly BUSCO summaries for representative assemblies of the three newly assembled genomes.
  - Supplementary Table S4. Master species table for all 17 sampled genomes.
  - Supplementary Table S5. Repeat composition summaries for the three newly assembled genomes.
  - Supplementary Table S6. Annotation BUSCO summaries for the primary annotations of the three newly assembled genomes.
  - Supplementary Table S7. Protein counts and BUSCO completeness for the harmonized 17-species annotation set.
  - Supplementary Table S8. Progressive Cactus guide/reference configurations and CASTER outcome summary.
  - Supplementary Table S9. Key-node support summary for the full ASTRAL concordance tree.
  - Supplementary Table S10. Gene genealogy interrogation (GGI) AU-test summary across the three constrained topologies.
  - Supplementary Table S11. Annotation-provenance sensitivity dataset assignments.
  - Supplementary Table S12. Whole-matrix microsynteny AU-test summary across the three constrained topology hypotheses.
  - Supplementary Table S13. Topology-informative microsynteny cluster summary.
  - Supplementary Table S14. Gene-adjacency maximum-likelihood and AU-test summary.
  - Supplementary Table S15. AGORA ancestral-reconstruction metric summary across workflow variants and topologies.
  - Supplementary Table S16. Dollo-like adjacency likelihood model-comparison summary.
  - Supplementary Table S17. DESCHRAMBLER resolution-selection summary across the original multi-resolution reference/topology runs.
  - Supplementary Table S18. Full-taxon DESCHRAMBLER run design and metric summary.
  - Supplementary Table S19. Full-taxon DESCHRAMBLER pairwise comparison statistics.
  - Supplementary Table S20. DESCHRAMBLER run-design and metric summary for the reduced same-depth rerun.
  - Supplementary Table S21. DESCHRAMBLER pairwise comparison statistics for the reduced same-depth rerun.
- Supplementary Figures
  - Supplementary Figure S1. k-mer distribution from GenomeScope2 for *Psettodes erumei*.
  - Supplementary Figure S2. *Psettodes erumei* chromosome assembly contact map.
  - Supplementary Figure S3. Full ROADIES species tree.
  - Supplementary Figure S4. Progressive Cactus to CASTER workflow summary.
  - Supplementary Figure S5. Reference-bias assessment for raw CASTER-pair trees.
  - Supplementary Figure S6. Reference-bias assessment for raw CASTER-site trees.
  - Supplementary Figure S7. Robinson-Foulds distance heatmap for raw CASTER-pair trees versus the two guide trees.
  - Supplementary Figure S8. Robinson-Foulds distance heatmap for raw CASTER-site trees versus the two guide trees.
  - Supplementary Figure S9. Robinson-Foulds distance heatmap comparing the post-processed CASTER trees to the guide trees.
  - Supplementary Figure S10. Pairwise Robinson-Foulds distance heatmap among the post-processed CASTER trees.
  - Supplementary Figure S11. Full ASTRAL concordance-factor tree.
  - Supplementary Figure S12. AU-ranking curves across three constrained gene-tree topologies used in the per-gene approximately unbiased tests.
  - Supplementary Figure S13. RefSeq/TOGA annotation-provenance microsynteny tree.
  - Supplementary Figure S14. BRAKER/GALBA annotation-provenance microsynteny tree.
  - Supplementary Figure S15. Microsynteny analysis workflow summary.
  - Supplementary Figure S16. Lower-triangular heatmap of pairwise microsynteny-conservation percentages among the 17 sampled genomes.
  - Supplementary Figure S17. Variant-only clustered microsynteny-profile heatmap summarizing synteny-cluster copy-number variation across the 17 sampled genomes.
  - Supplementary Figure S18. Full-species microsynteny tree inferred from MCScanX collinearity with syntenet.
  - Supplementary Figure S19. Whole-matrix microsynteny AU-test summary across the three constrained topology hypotheses.
  - Supplementary Figure S20. Annotated heatmap of normalized pairwise adjacency distances for the primary strict full-species microsynteny dataset.
  - Supplementary Figure S21. Complete Minimum Evolution tree from the primary strict full-species microsynteny distance matrix.
  - Supplementary Figure S22. Annotated heatmap of normalized pairwise adjacency distances for the no-*Solea* microsynteny sensitivity dataset.
  - Supplementary Figure S23. Complete Minimum Evolution tree from the no-*Solea* microsynteny distance matrix.
  - Supplementary Figure S24. Binary adjacency maximum-likelihood trees from the all-scaffolds and scaffold-filtered datasets.
  - Supplementary Figure S25. Binary adjacency AU-test summary across the three constrained topology hypotheses.
  - Supplementary Figure S26. Macrosynteny analysis workflow summary.
  - Supplementary Figure S27. Macrosynteny chromosome-comparison oxford dot plots for *Psettodes erumei*.
  - Supplementary Figure S28. Macrosynteny ribbon plots under the three alternative topology hypotheses.
  - Supplementary Figure S29. Quartet genome rearrangement simulation summary for FM-compatible taxon sets.
  - Supplementary Figure S30. Quartet genome rearrangement simulation summary for FP1-compatible taxon sets.
  - Supplementary Figure S31. Quartet genome rearrangement simulation summary for FP2-compatible taxon sets.
  - Supplementary Figure S32. Quintet genome rearrangement simulation summary for FM-compatible taxon sets.
  - Supplementary Figure S33. Quintet genome rearrangement simulation summary for FP2-compatible taxon sets.
  - Supplementary Figure S34. DESCHRAMBLER design-shift summary from the all-tips design to the reduced same-depth design.
  - Supplementary Figure S35. DESCHRAMBLER evidence matrix for the all-tips FM vs FP1 comparison.
  - Supplementary Figure S36. DESCHRAMBLER evidence matrix for the all-tips FM vs FP2 comparison.
  - Supplementary Figure S37. DESCHRAMBLER evidence matrix for the reduced same-depth FM vs FP1 comparison.
  - Supplementary Figure S38. DESCHRAMBLER evidence matrix for the reduced same-depth FM vs FP2 comparison.

### Supplementary Tables

##### **Supplementary Table S1.** Specimen, voucher, sequencing-library, and accession metadata for the three newly assembled genomes.

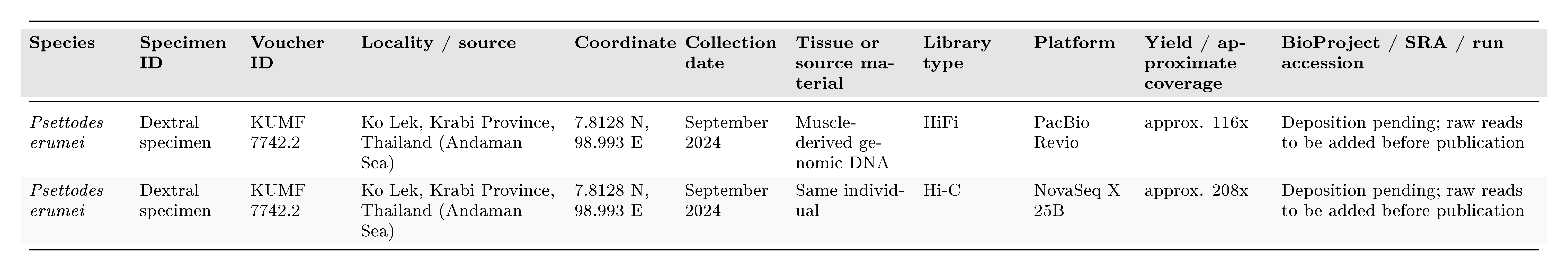

##### **Supplementary Table S2.** Genome assembly statistics for representative assemblies of the three newly assembled genomes.

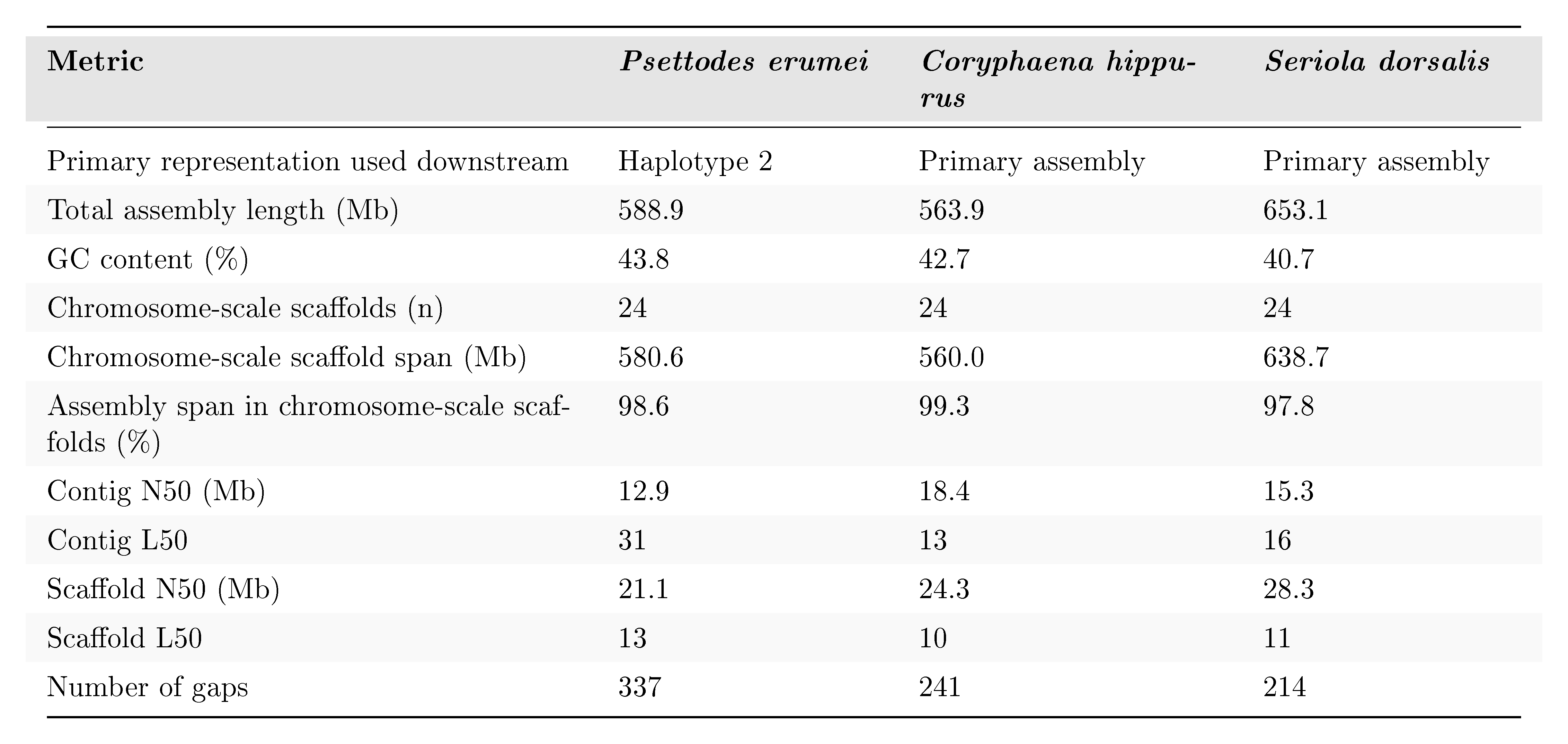

##### **Supplementary Table S3.** Assembly BUSCO summaries for representative assemblies of the three newly assembled genomes.

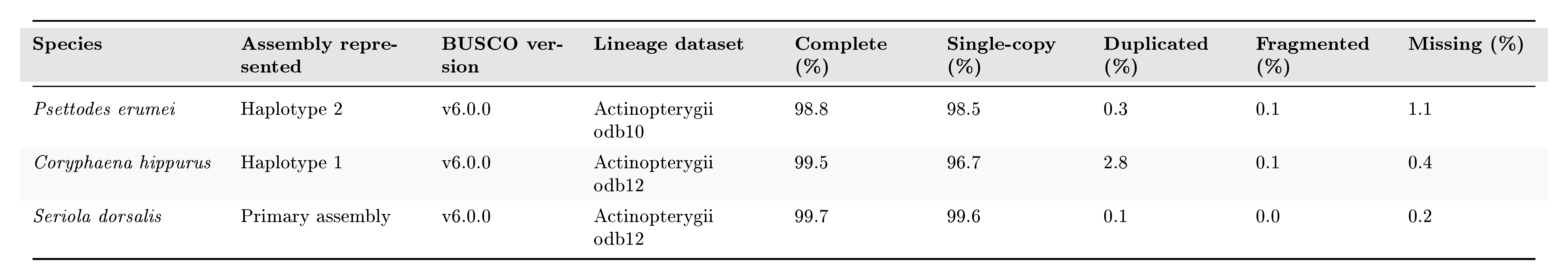

##### **Supplementary Table S4.** Master species table for all 17 sampled genomes.

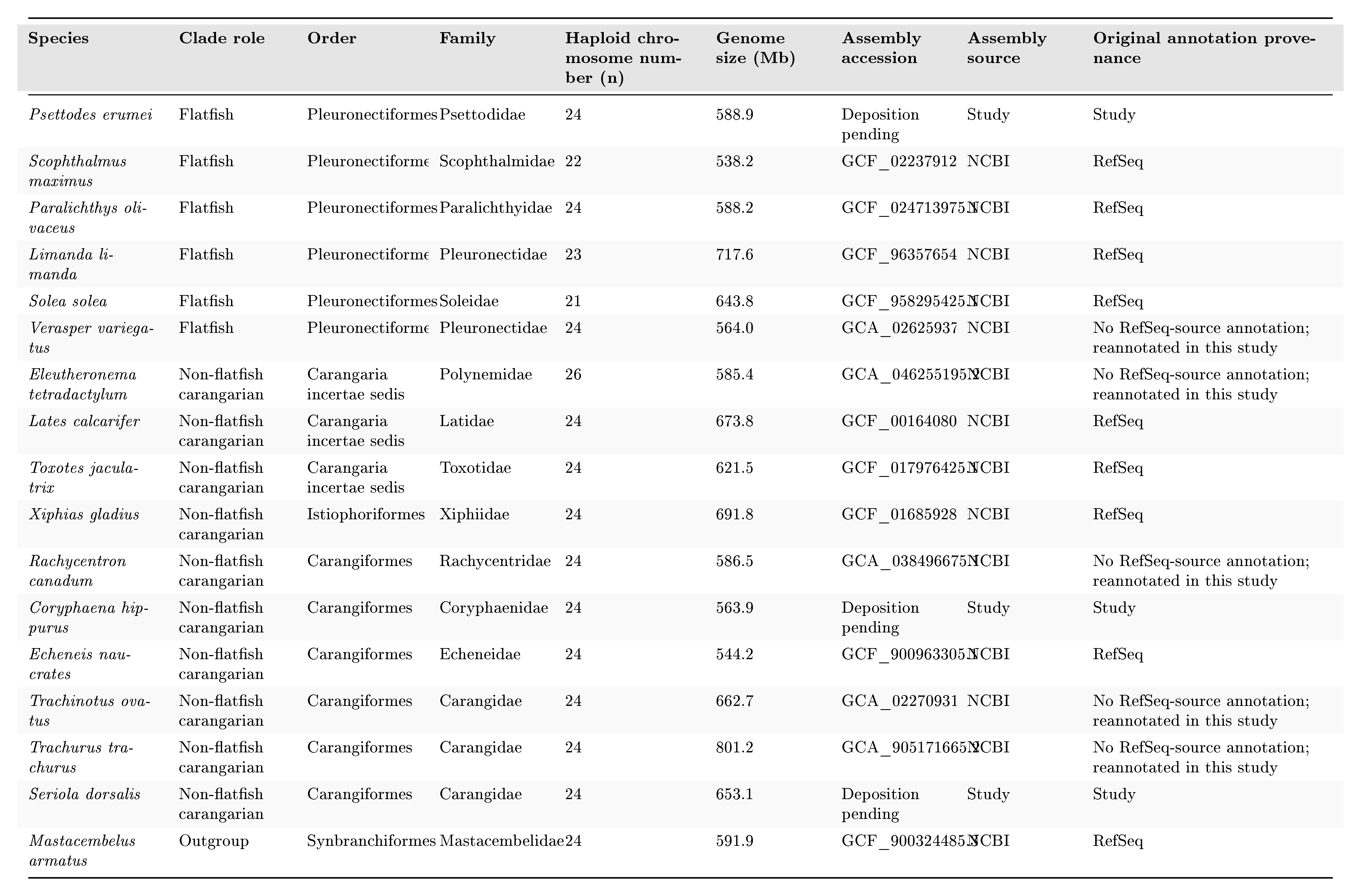

##### **Supplementary Table S5.** Repeat composition summaries for the three newly assembled genomes.

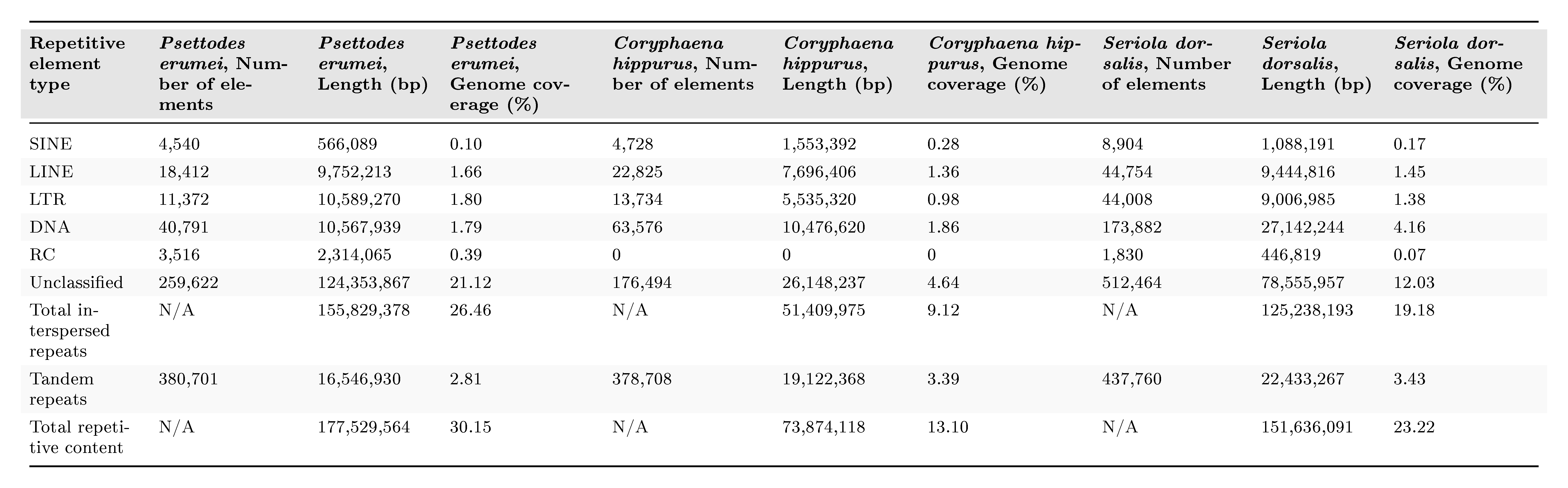

##### **Supplementary Table S6.** Annotation BUSCO summaries for the primary annotations of the three newly assembled genomes.

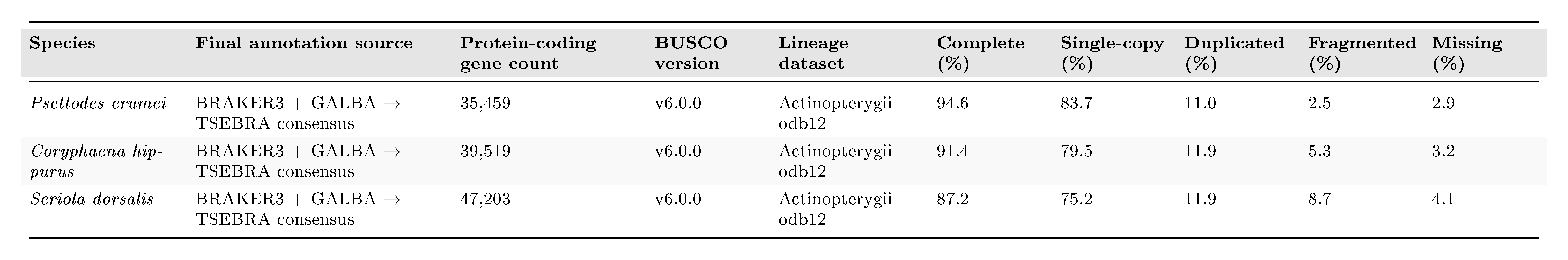

##### **Supplementary Table S7.** Protein counts and BUSCO completeness for the harmonized 17-species annotation set.

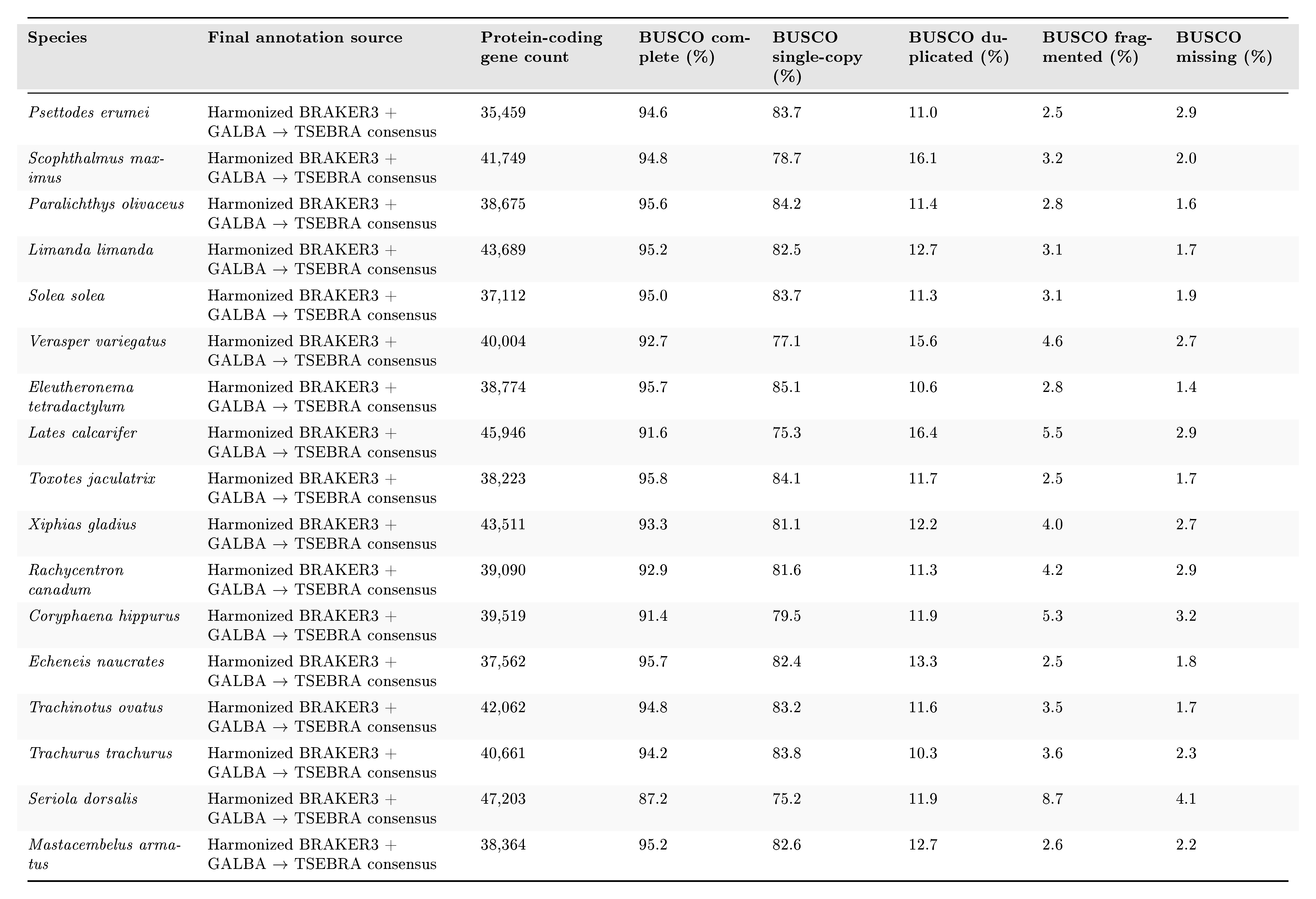

##### **Supplementary Table S8.** Progressive Cactus guide/reference configurations and CASTER outcome summary.

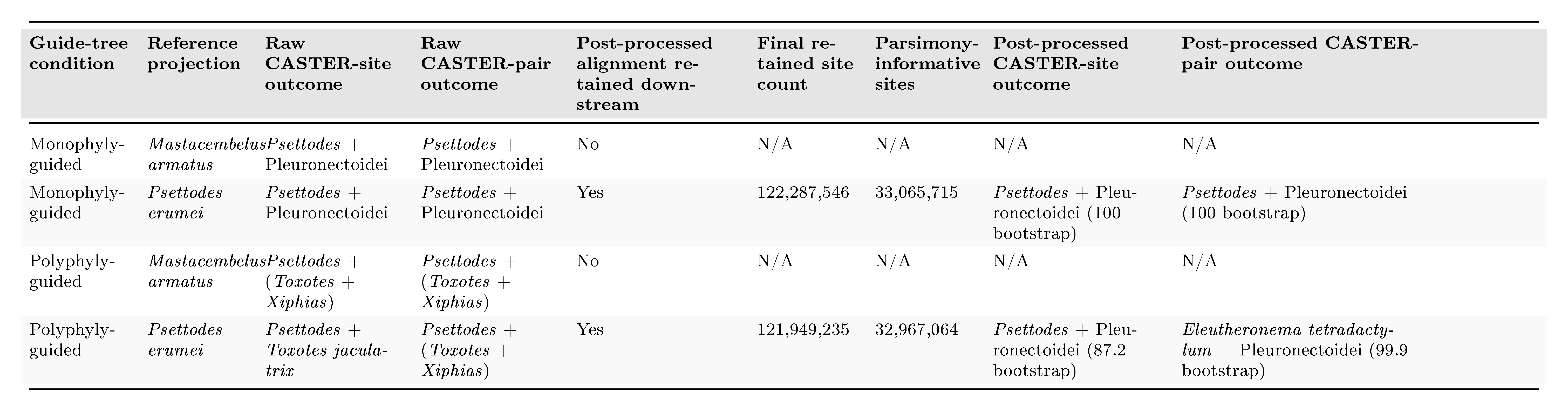

##### **Supplementary Table S9.** Key-node support summary for the full ASTRAL concordance tree.

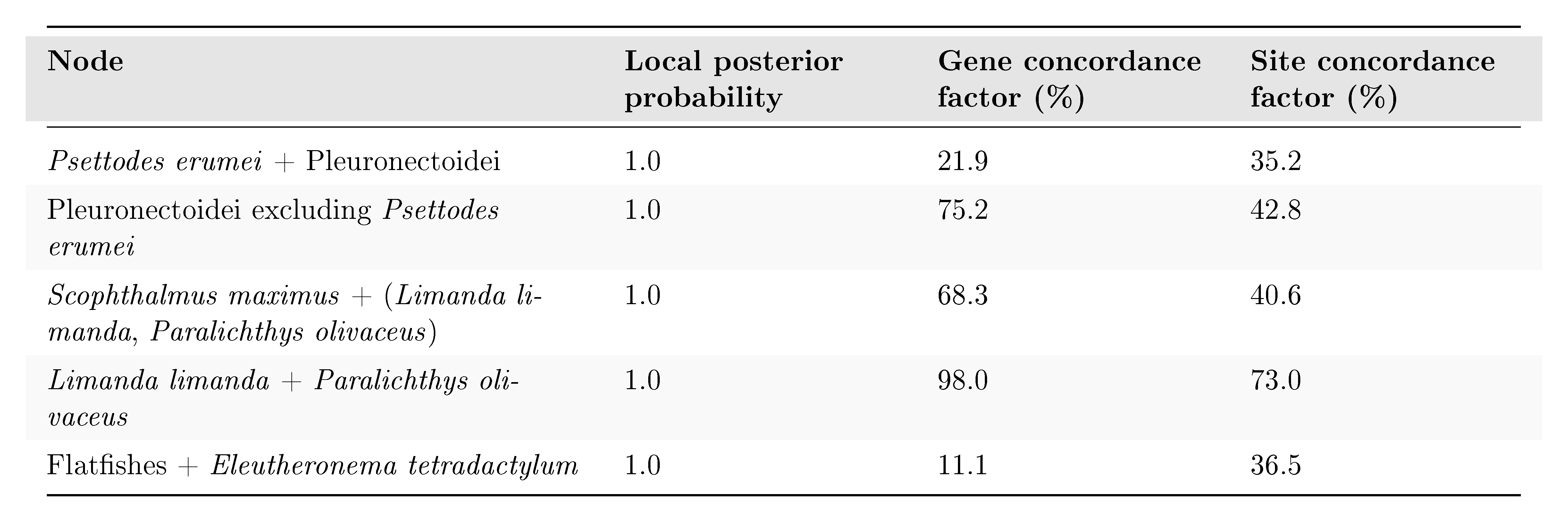

##### **Supplementary Table S10.** Gene genealogy interrogation (GGI) AU-test summary across the three constrained topologies.

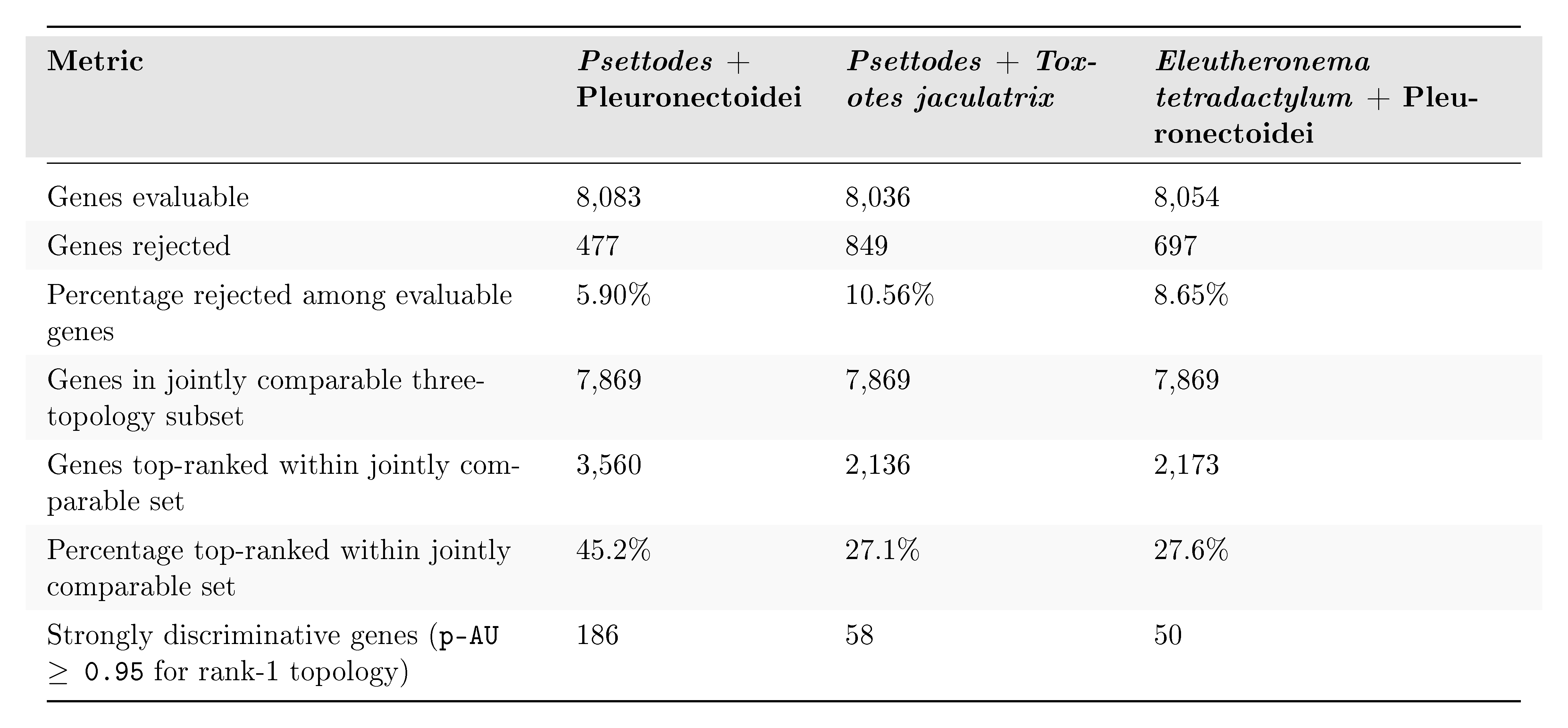

##### **Supplementary Table S11.** Annotation-provenance sensitivity dataset assignments.

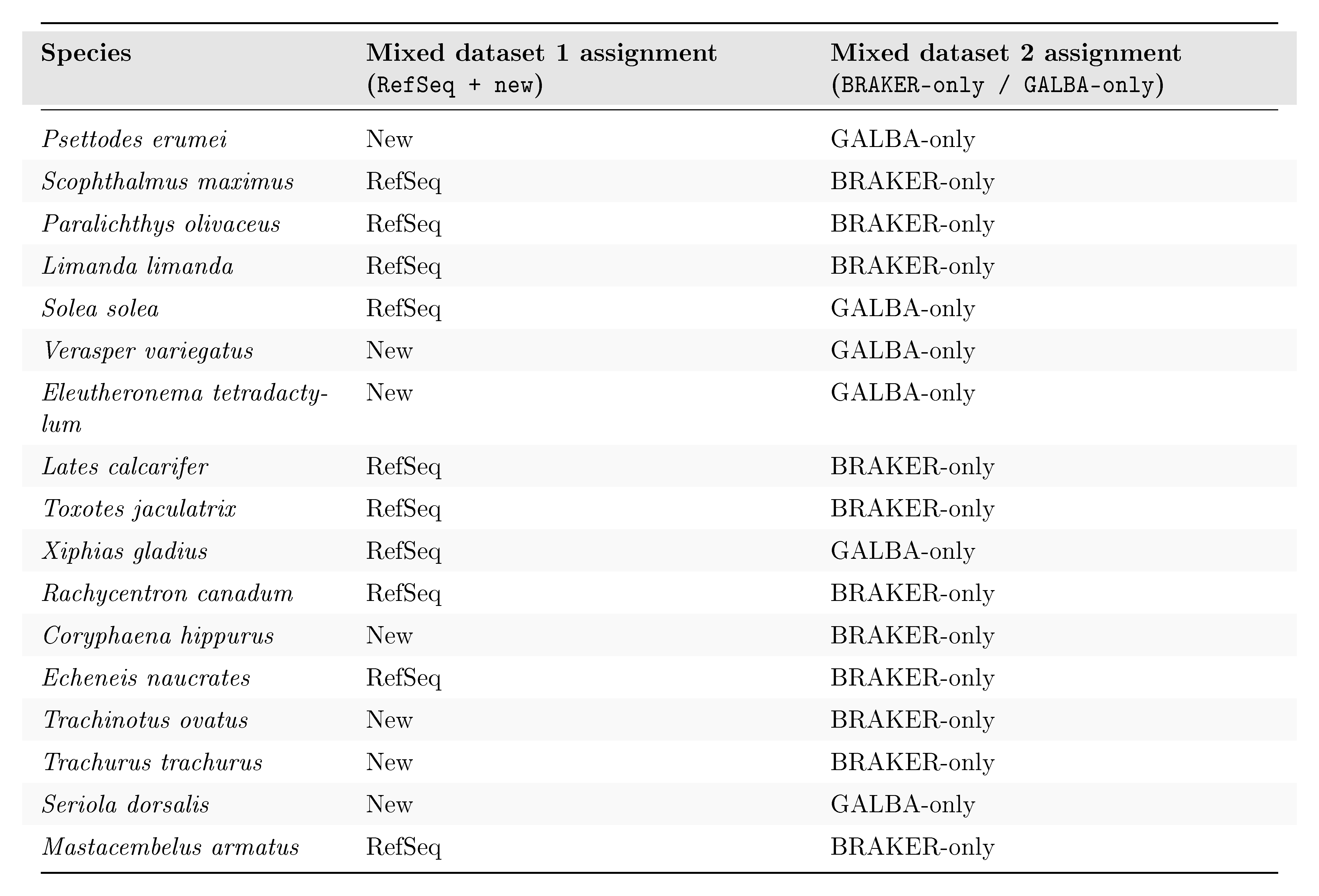

##### **Supplementary Table S12.** Whole-matrix microsynteny AU-test summary across the three constrained topology hypotheses.

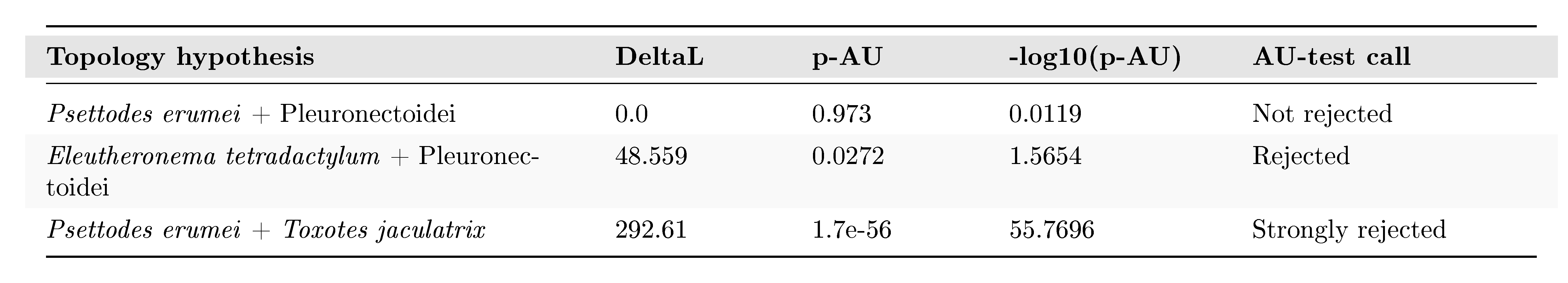

##### **Supplementary Table S13.** Topology-informative microsynteny cluster summary.

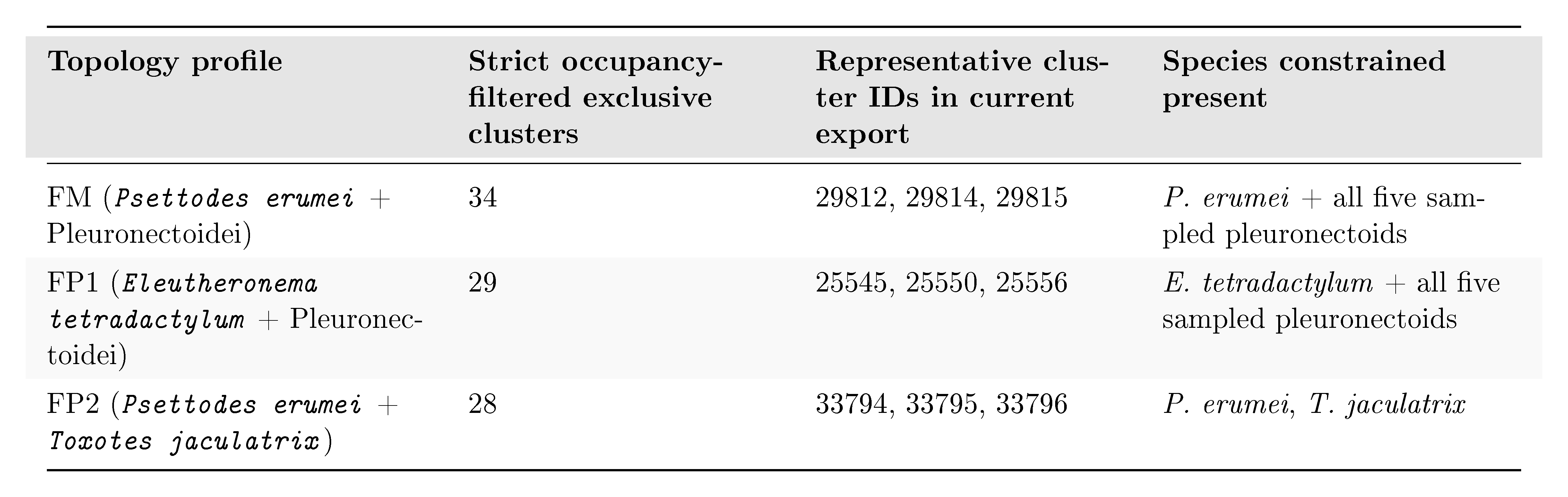

##### **Supplementary Table S14.** Gene-adjacency maximum-likelihood and AU-test summary.

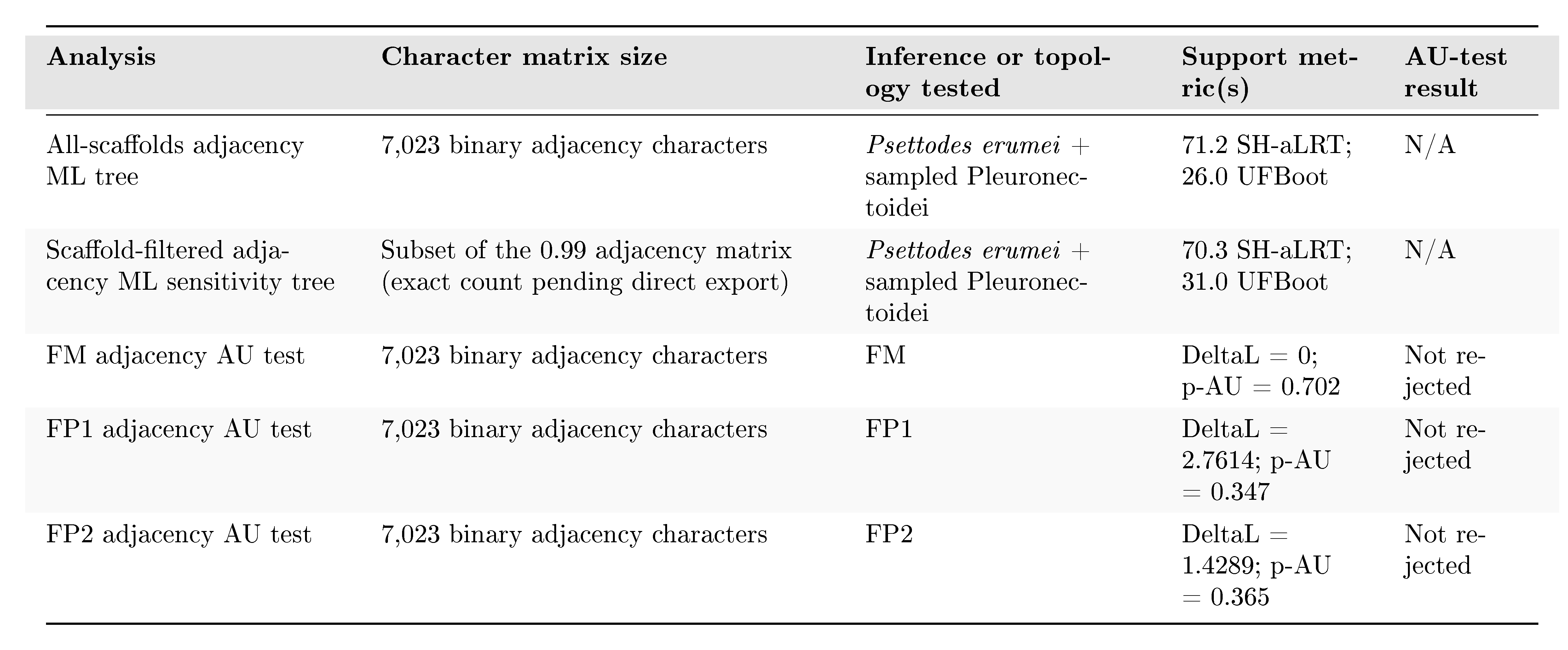

##### **Supplementary Table S15.** AGORA ancestral-reconstruction metric summary across workflow variants and topologies.

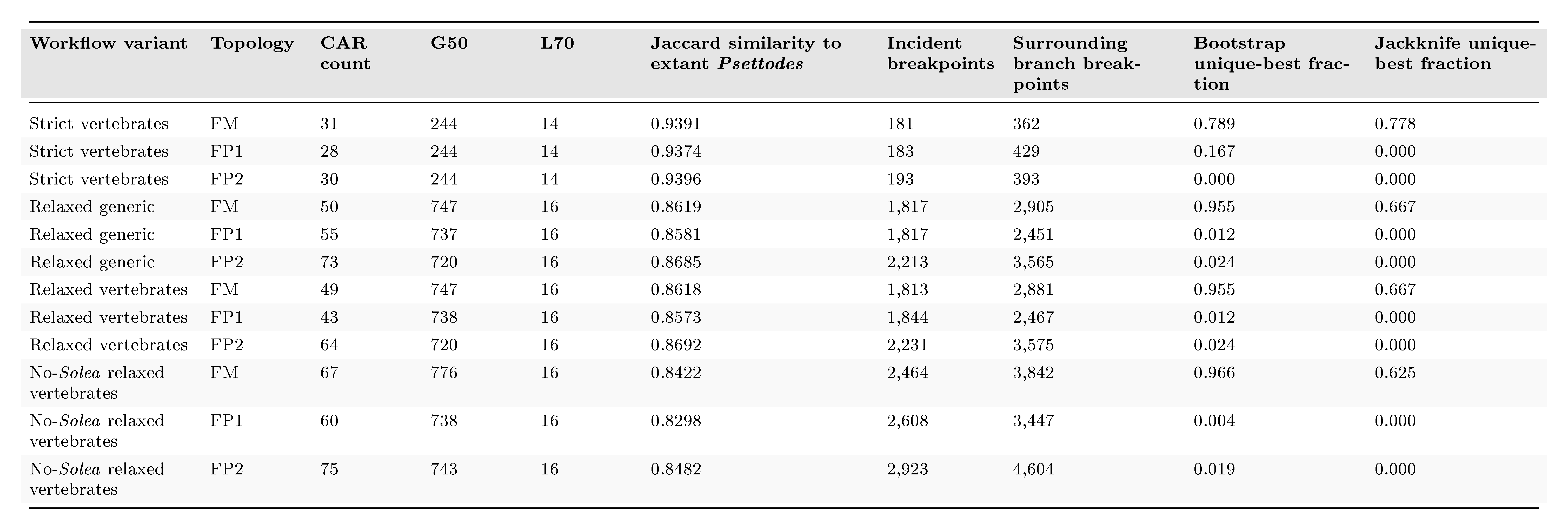

##### **Supplementary Table S16.** Dollo-like adjacency likelihood model-comparison summary.

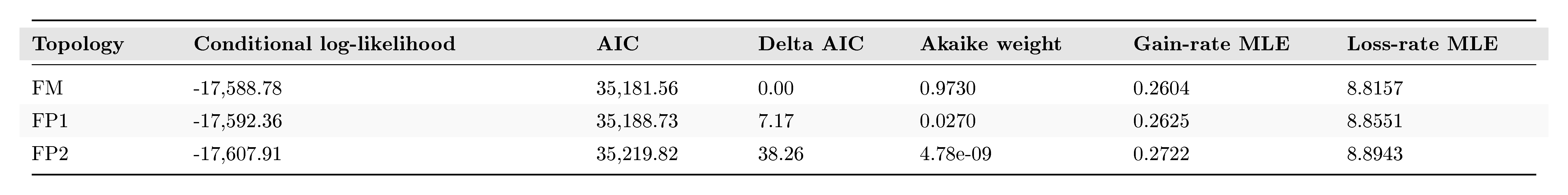

##### **Supplementary Table S17.** DESCHRAMBLER resolution-selection summary across the original multi-resolution reference/topology runs.

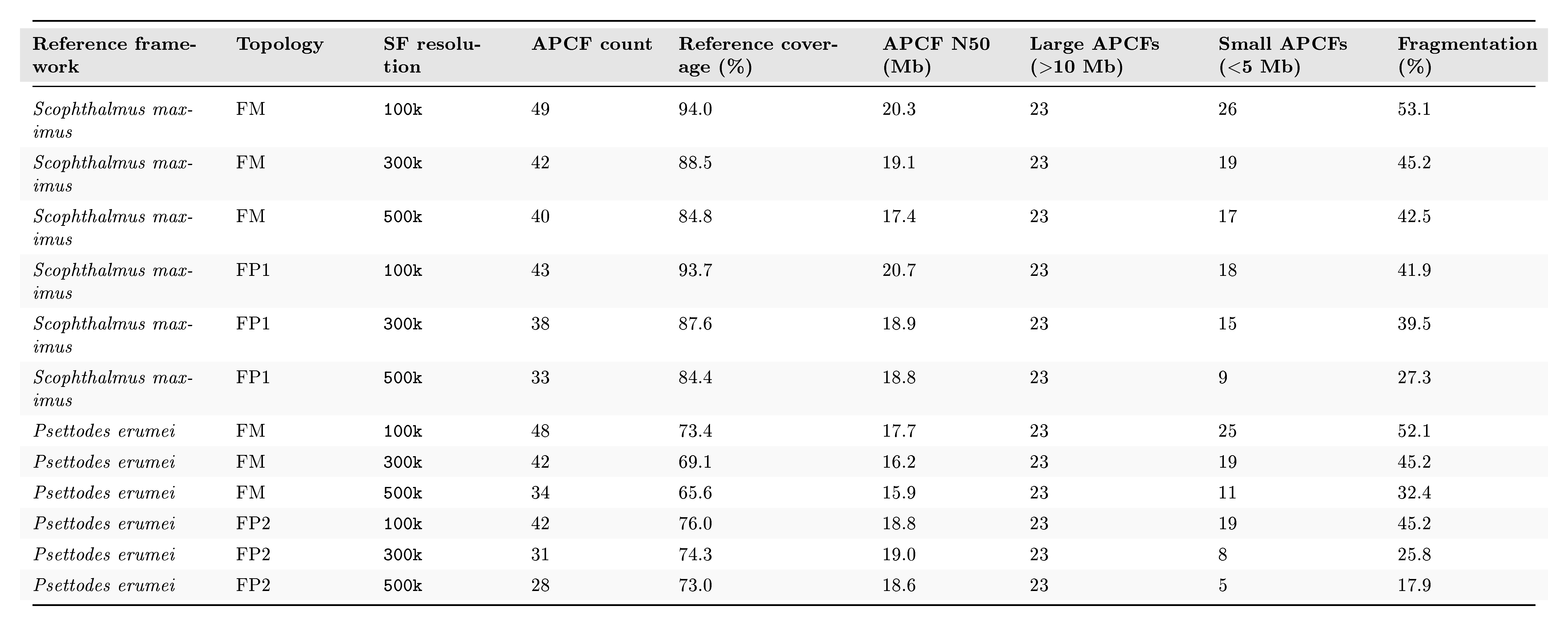

##### **Supplementary Table S18.** Full-taxon DESCHRAMBLER run design and metric summary.

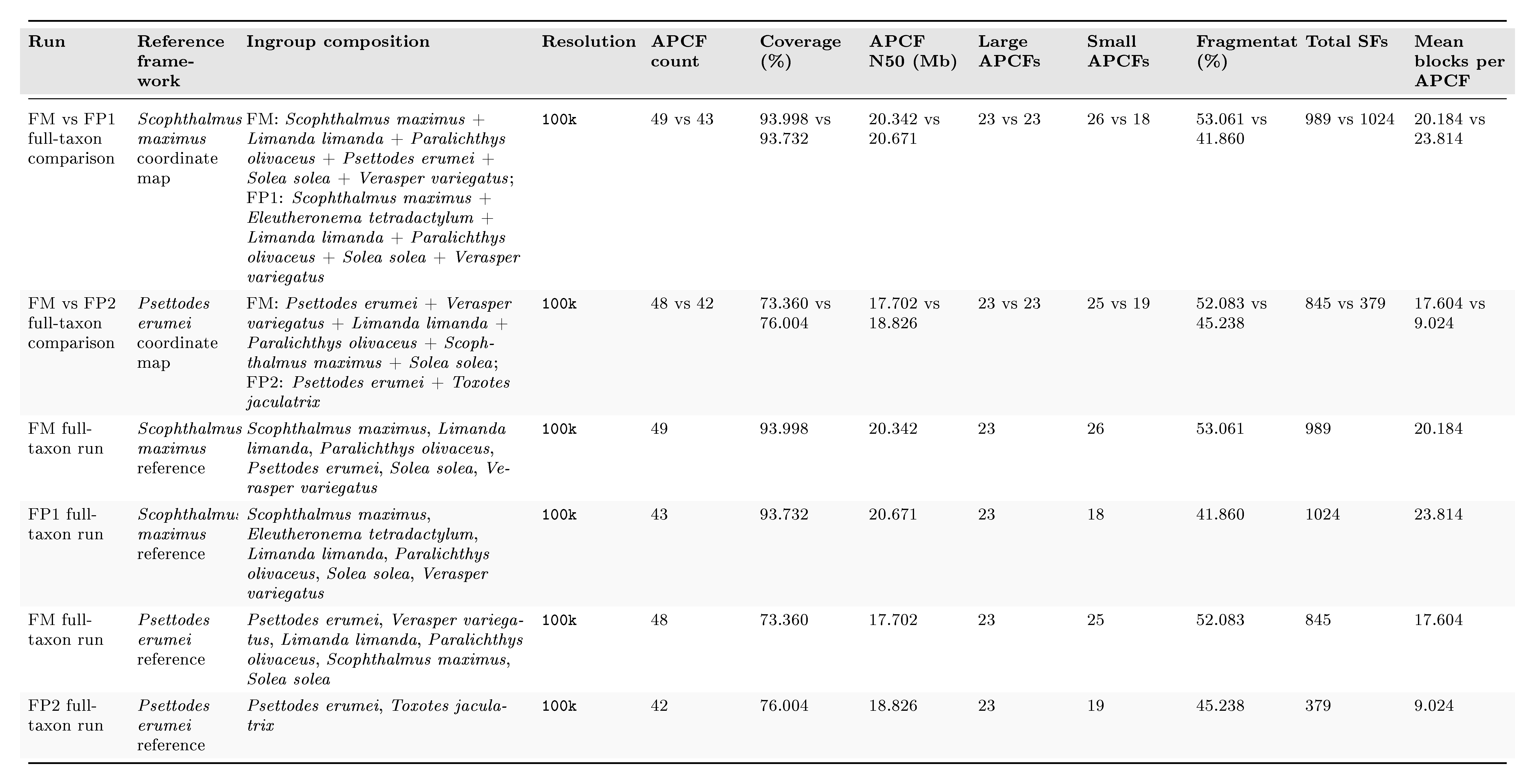

##### **Supplementary Table S19.** Full-taxon DESCHRAMBLER pairwise comparison statistics.

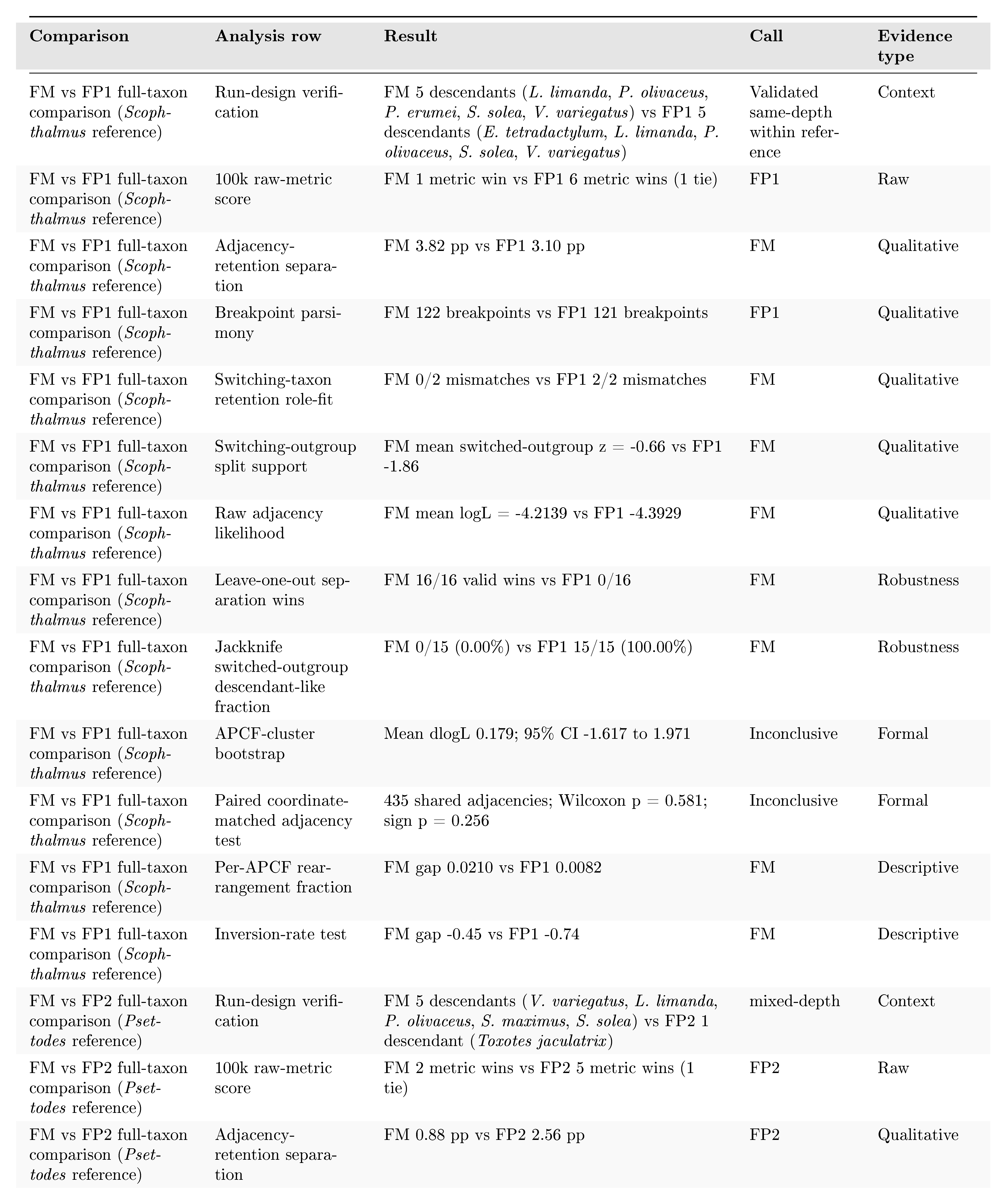

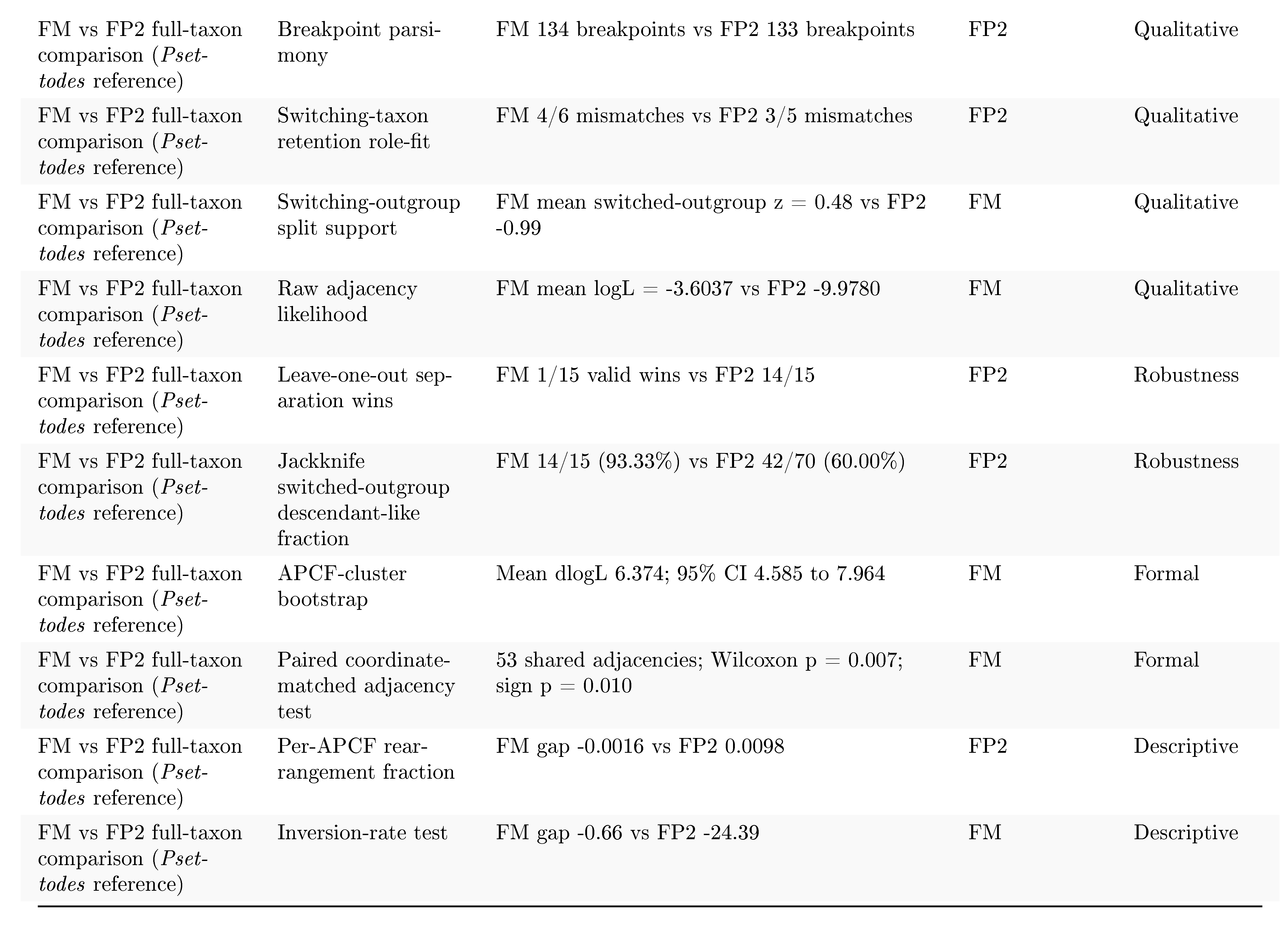

##### **Supplementary Table S20.** DESCHRAMBLER run-design and metric summary for the reduced same-depth rerun.

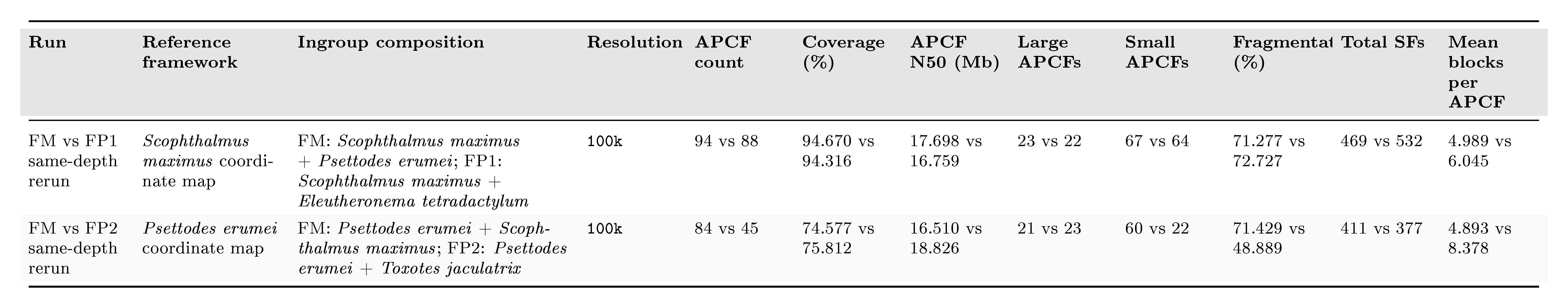

##### **Supplementary Table S21.** DESCHRAMBLER pairwise comparison statistics for the reduced same-depth rerun.

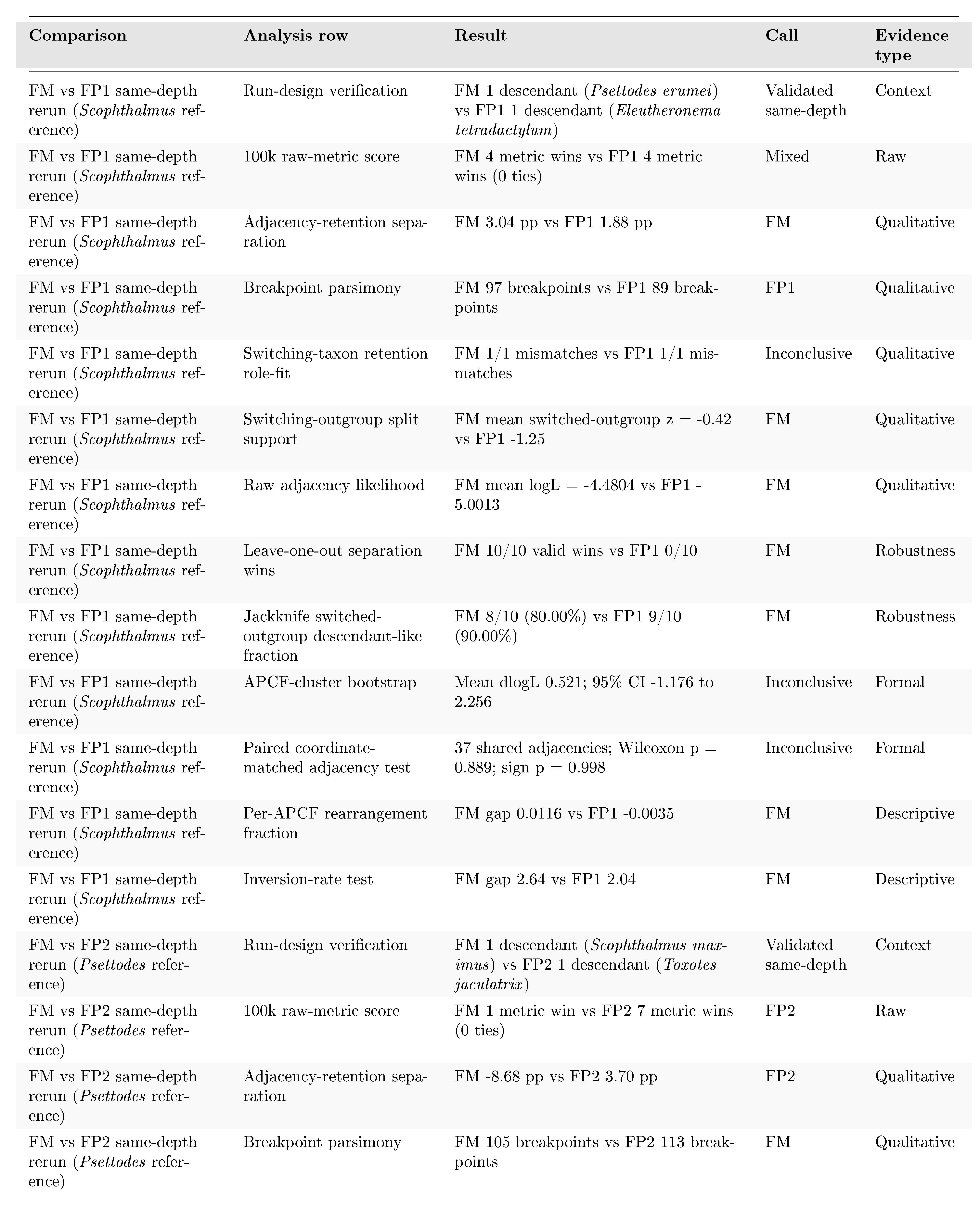

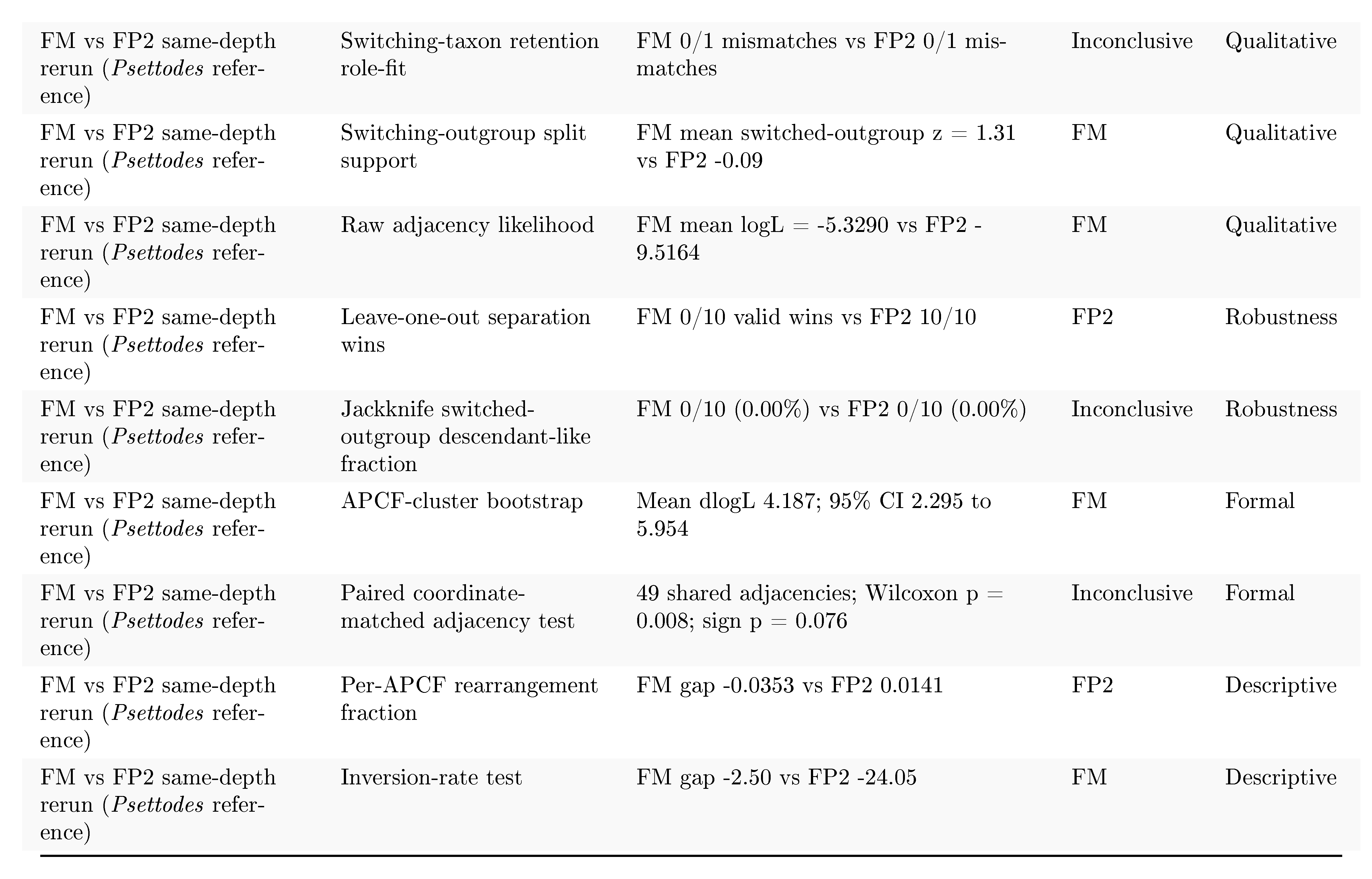

### Supplementary Figures

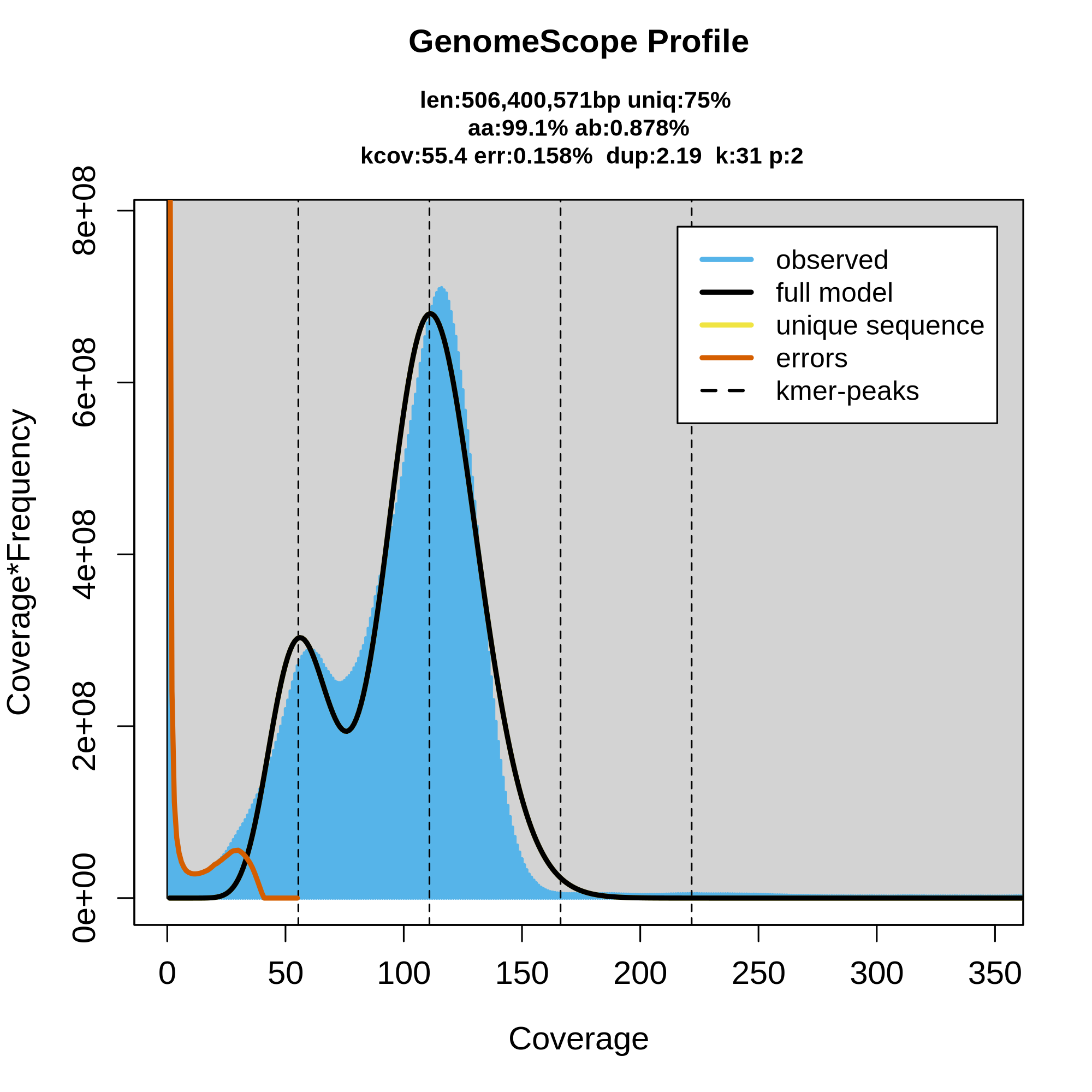

**Supplementary Figure S1.** k-mer distribution from GenomeScope2 for *Psettodes erumei*. GenomeScope2-style transformed k-mer distribution used to summarize the genome-size and repeat-profile signal for *Psettodes erumei*.

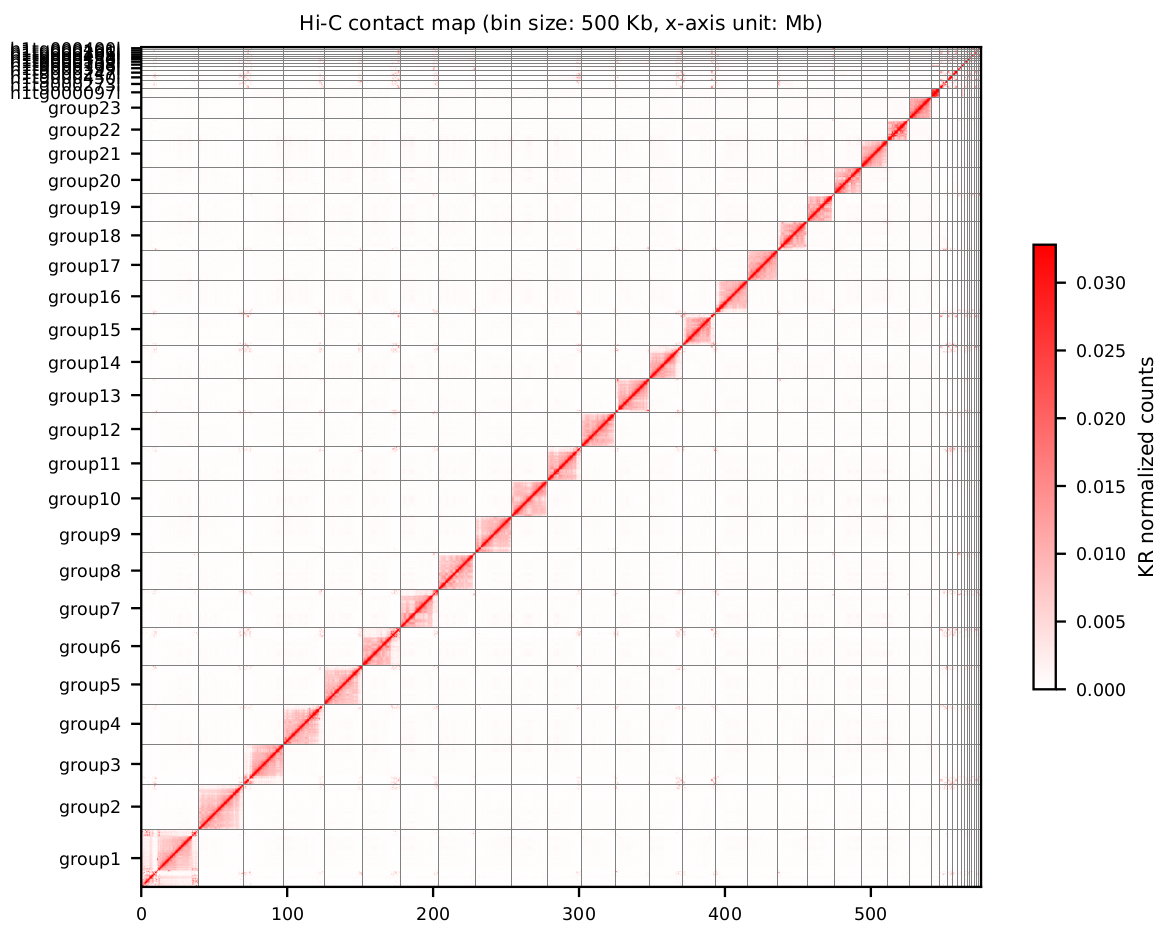

**Supplementary Figure S2.** *Psettodes erumei* chromosome assembly contact map. Hi-C contact heat map demonstrating the structural integrity of the *Psettodes erumei* chromosome-level assembly. The prominent diagonal squares indicate strong intra-chromosomal interactions, successfully anchoring and orienting the contigs into 24 distinct chromosomes.

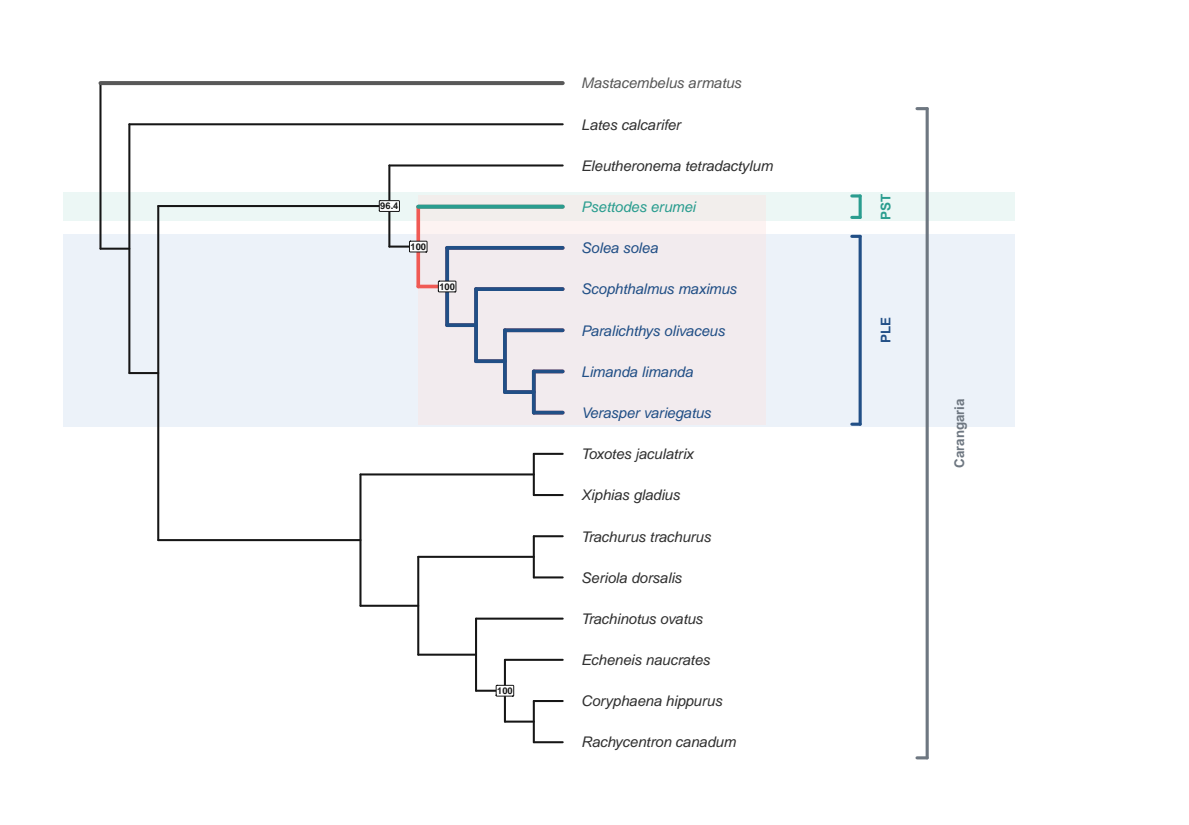

**Supplementary Figure S3.** Full ROADIES species tree. Full ROADIES / ASTRAL-Pro2 species tree for the 17-species dataset, rooted with *Mastacembelus armatus*. The inferred topology places *Psettodes erumei* as sister to the sampled Pleuronectoidei, with branch support values shown at internal nodes.

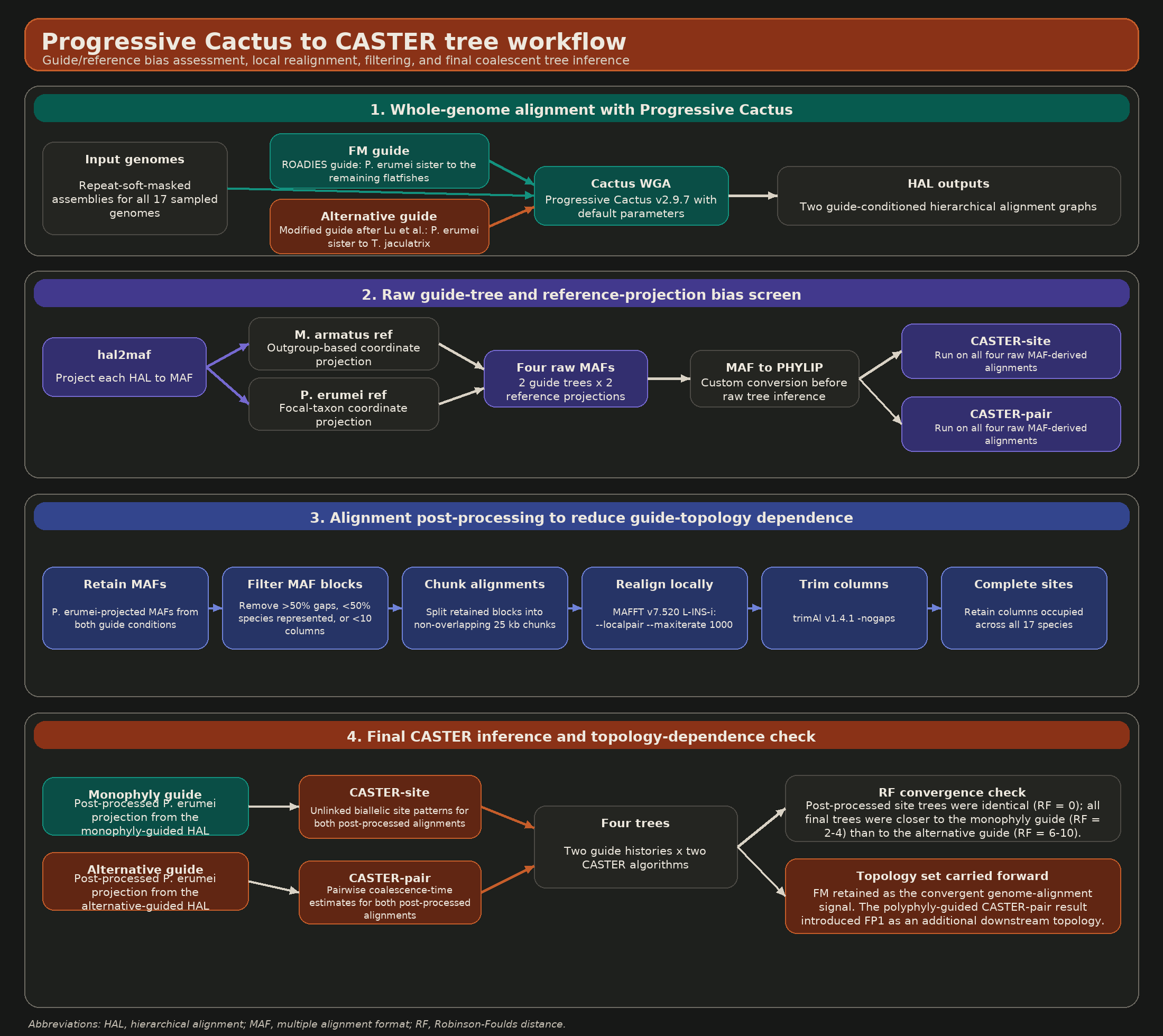

**Supplementary Figure S4.** Progressive Cactus to CASTER workflow summary. Flowchart summarizing the whole-genome alignment and CASTER species-tree workflow, from Progressive Cactus alignments through raw guide/reference-bias assessment, MAF filtering, 25-kb chunking, MAFFT realignment, trimAl gap-free site retention, and final CASTER-site/CASTER-pair species-tree inference with RF distance checks.

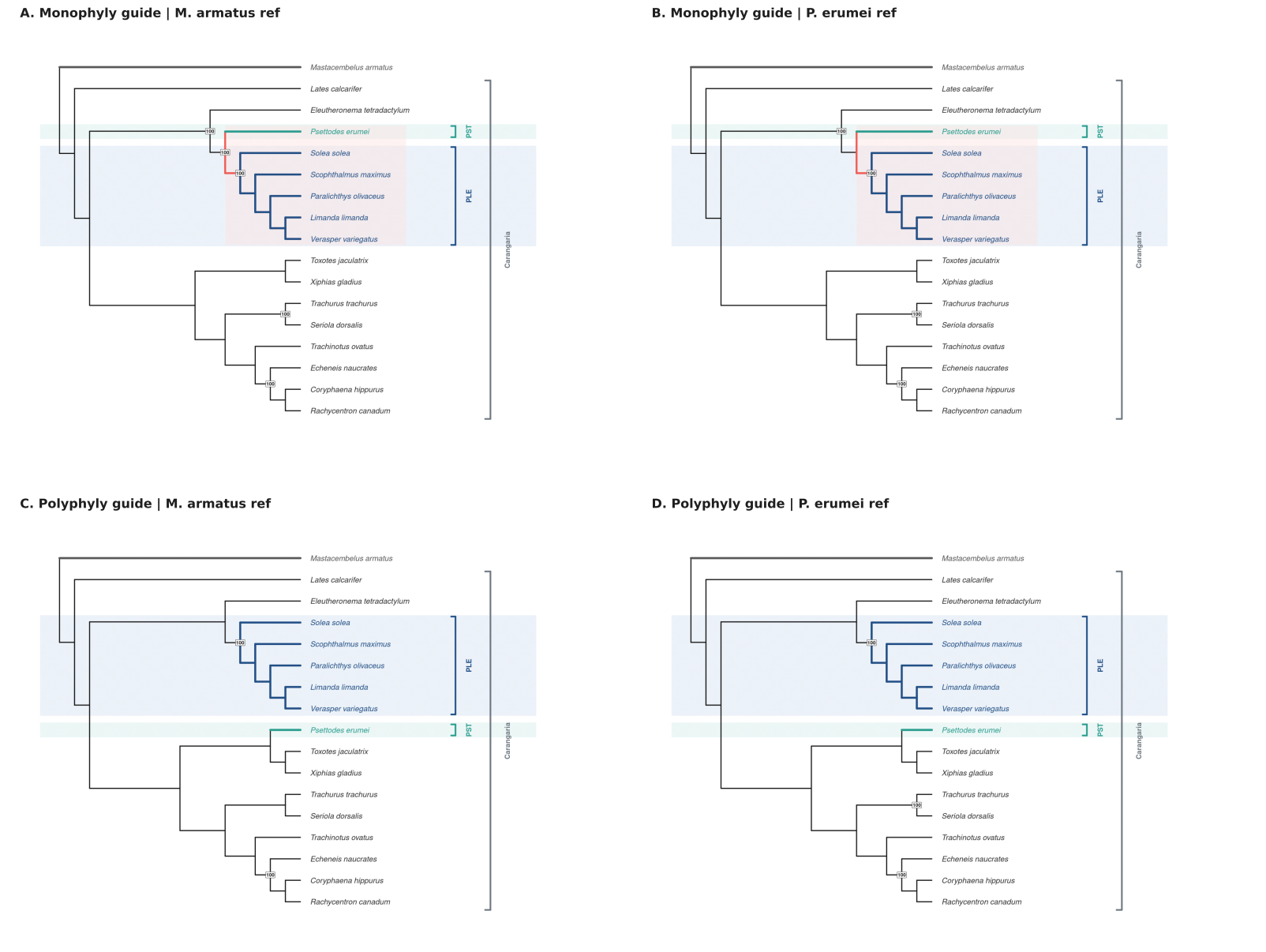

**Supplementary Figure S5.** Reference-bias assessment for raw CASTER-pair trees. Raw CASTER-pair trees from the four guide-tree and reference-projection combinations. The raw pair-based trees track the guide-tree condition closely, indicating strong alignment-conditioning bias before post-processing.

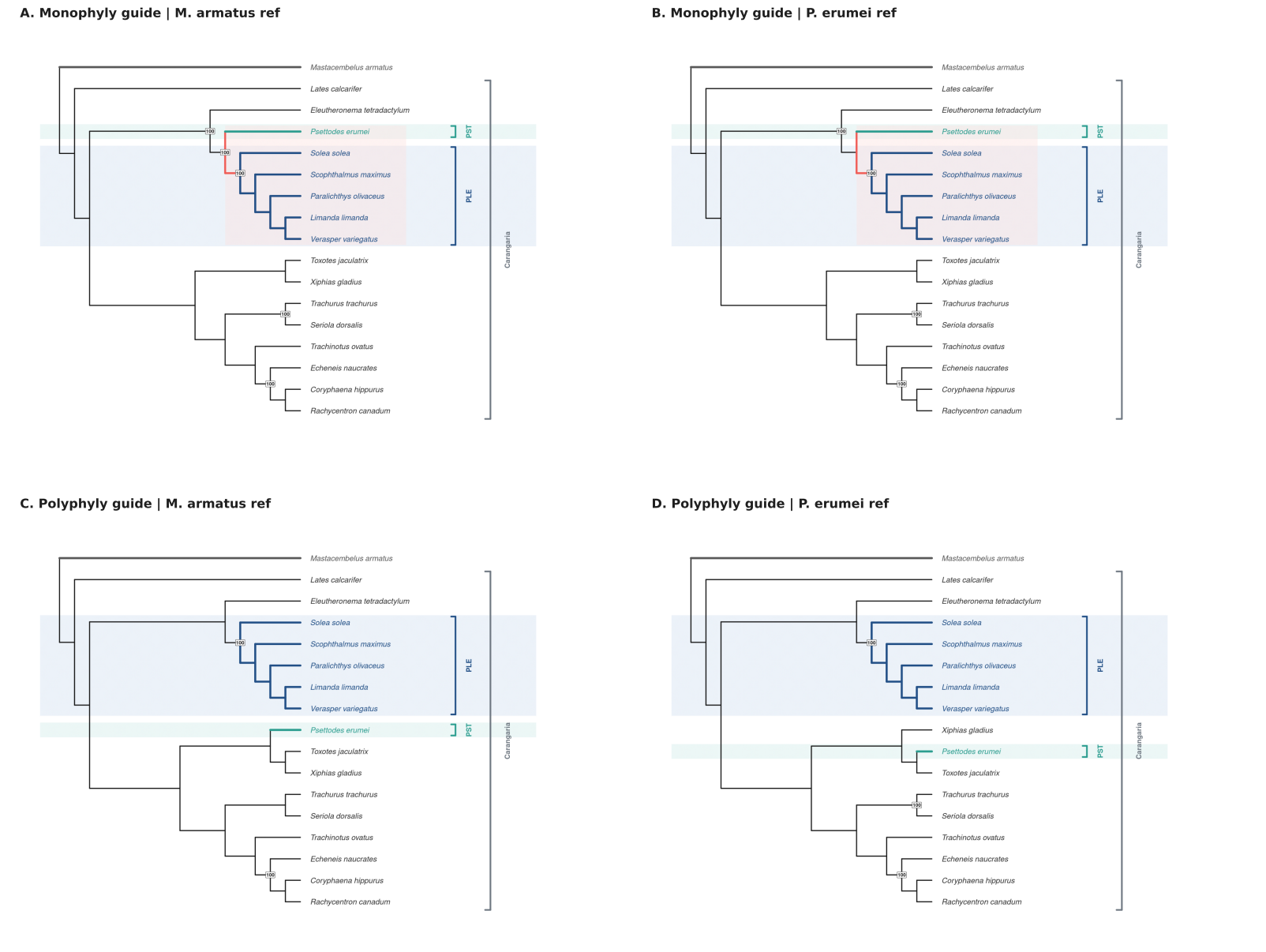

**Supplementary Figure S6.** Reference-bias assessment for raw CASTER-site trees. Raw CASTER-site trees from the same four guide-tree and reference-projection combinations. Site-based inference shows the same guide-tree dependence, with additional reference-projection sensitivity under the alternative guide.

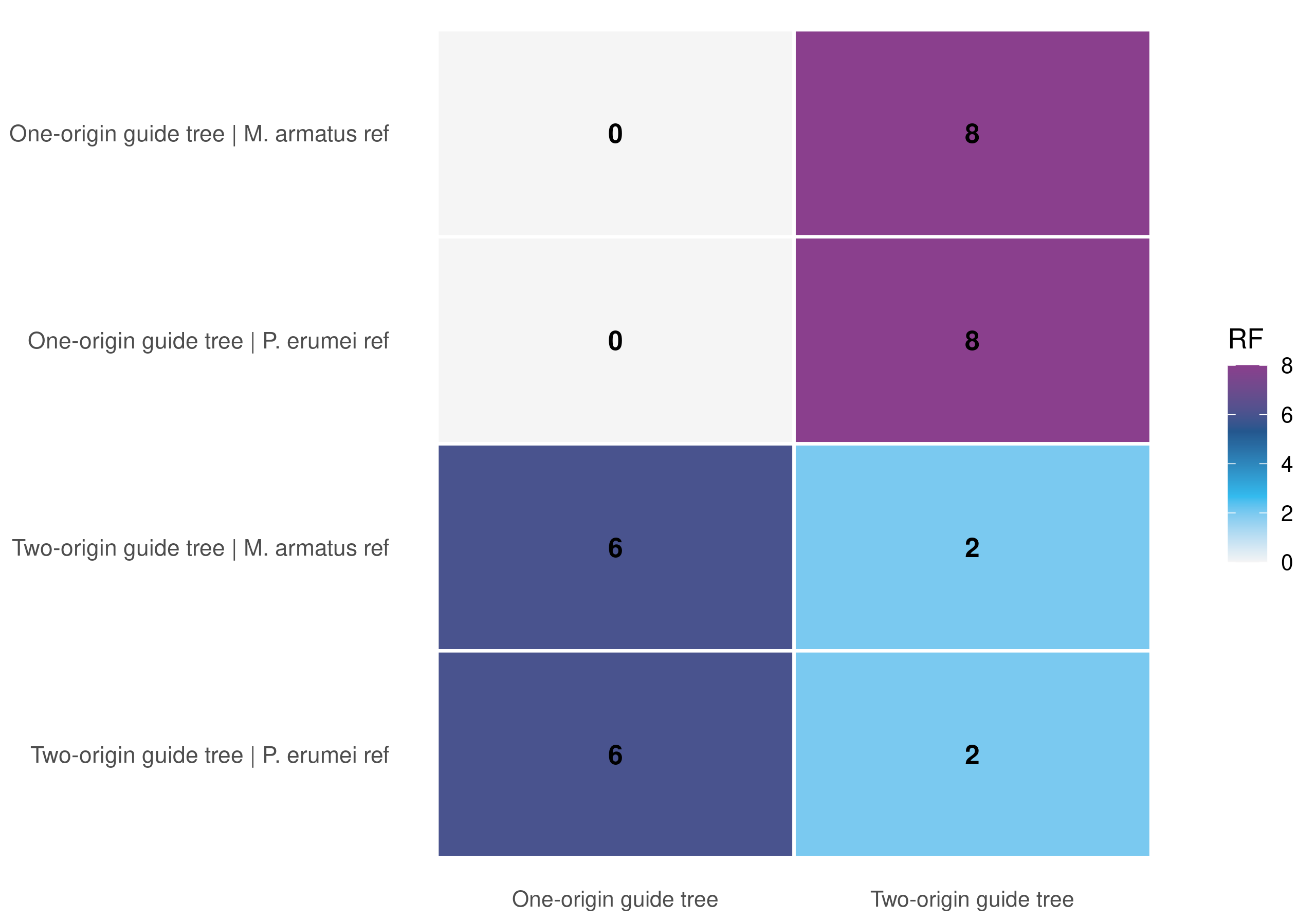

**Supplementary Figure S7.** Robinson-Foulds distance heatmap for raw CASTER-pair trees versus the two guide trees. Unrooted RF distances between the four raw CASTER-pair trees and the two alternative guide trees. The one-origin raw trees are identical to the monophyly guide, whereas the alternative-guided raw trees remain closer to the modified guide than to the monophyly guide.

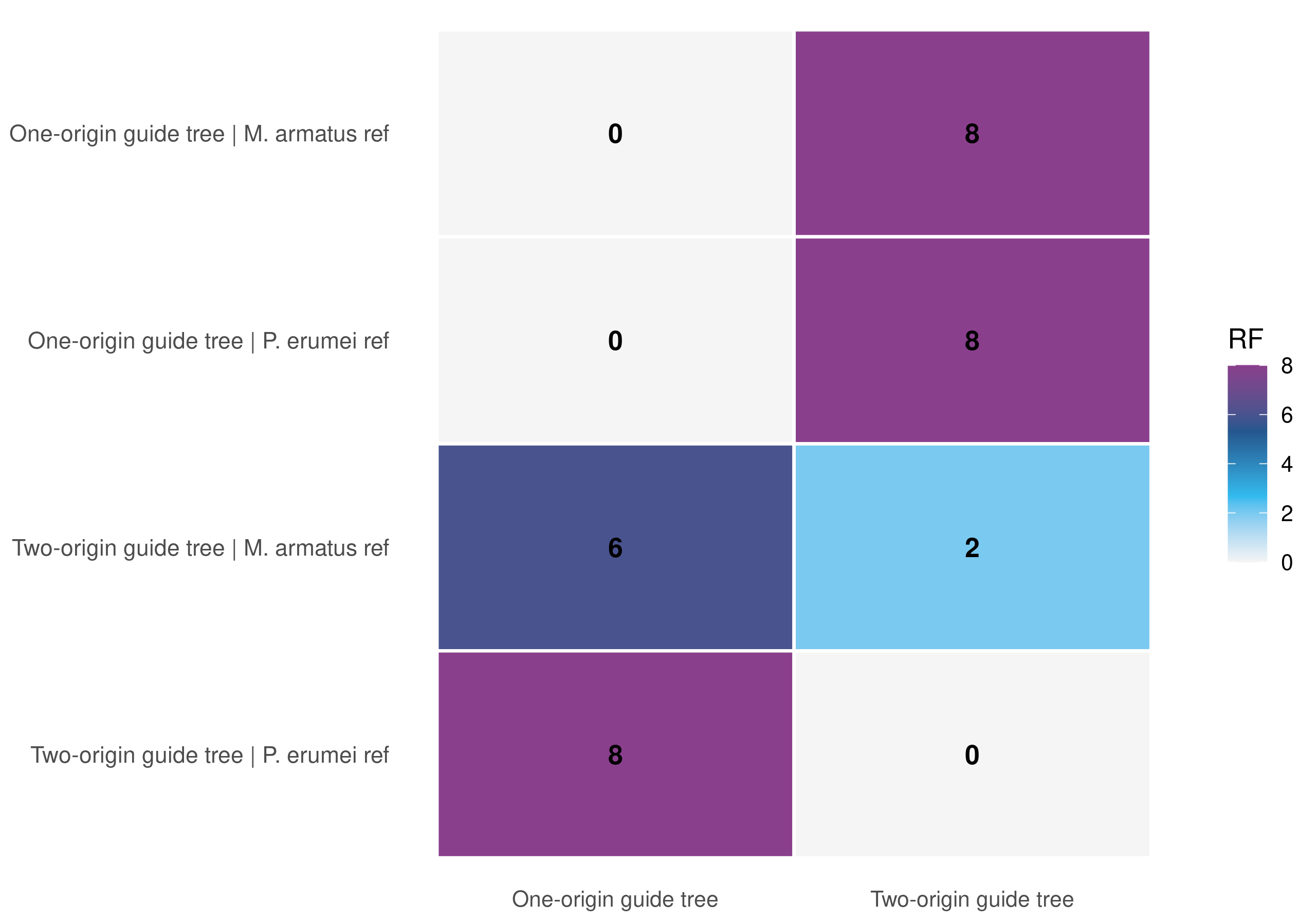

**Supplementary Figure S8.** Robinson-Foulds distance heatmap for raw CASTER-site trees versus the two guide trees. Unrooted RF distances between the four raw CASTER-site trees and the two alternative guide trees. These comparisons show the strongest match between each raw site tree and the guide condition under which its alignment was generated.

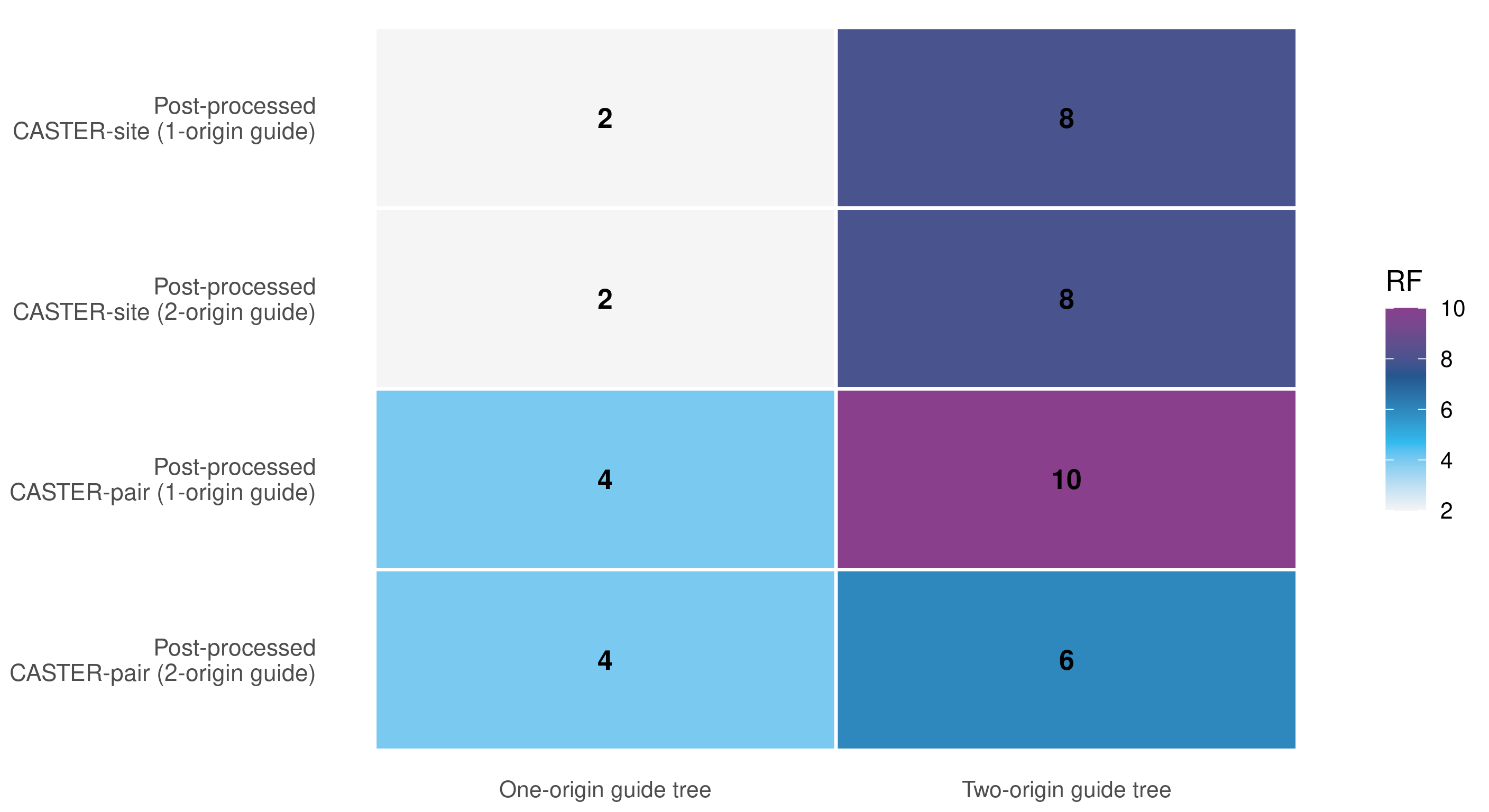

**Supplementary Figure S9.** Robinson-Foulds distance heatmap comparing the post-processed CASTER trees to the guide trees. Unrooted RF distances between the four post-processed CASTER trees and the two guide trees. The retained trees converge toward the monophyly-compatible solution rather than simply recapitulating the original guide topology.

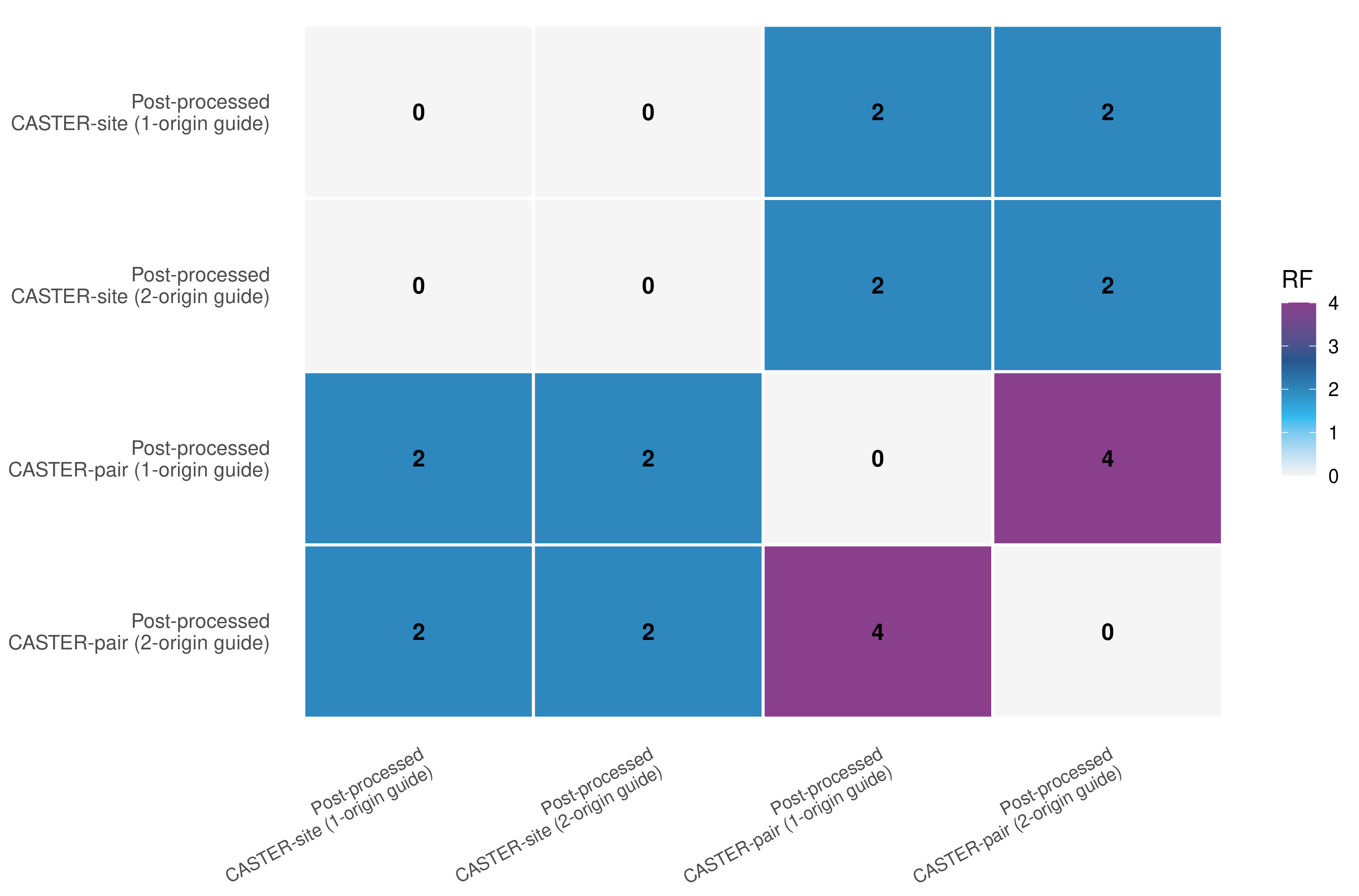

**Supplementary Figure S10.** Pairwise Robinson-Foulds distance heatmap among the post-processed CASTER trees. Pairwise RF distances among the four retained post-processed CASTER trees. The two site-based trees are identical, and the pair-based trees differ only modestly from that shared solution.

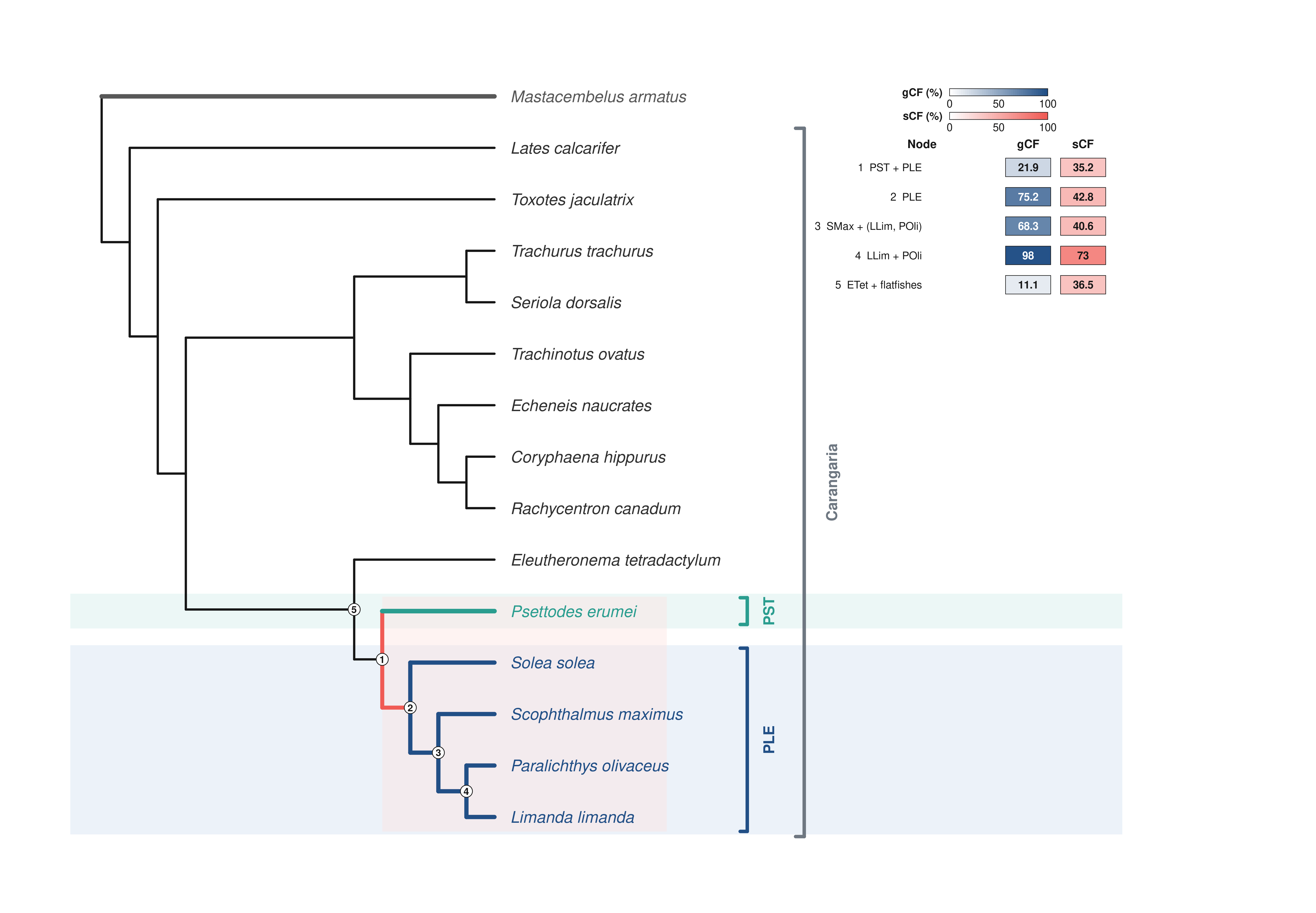

**Supplementary Figure S11.** Full ASTRAL concordance-factor tree. Full ASTRAL-III species tree inferred from 10,430 single-copy orthologs. Paired pies show gene concordance factor support on the left and site concordance factor support on the right, with the key nodes summarized numerically in Supplementary Table S9.

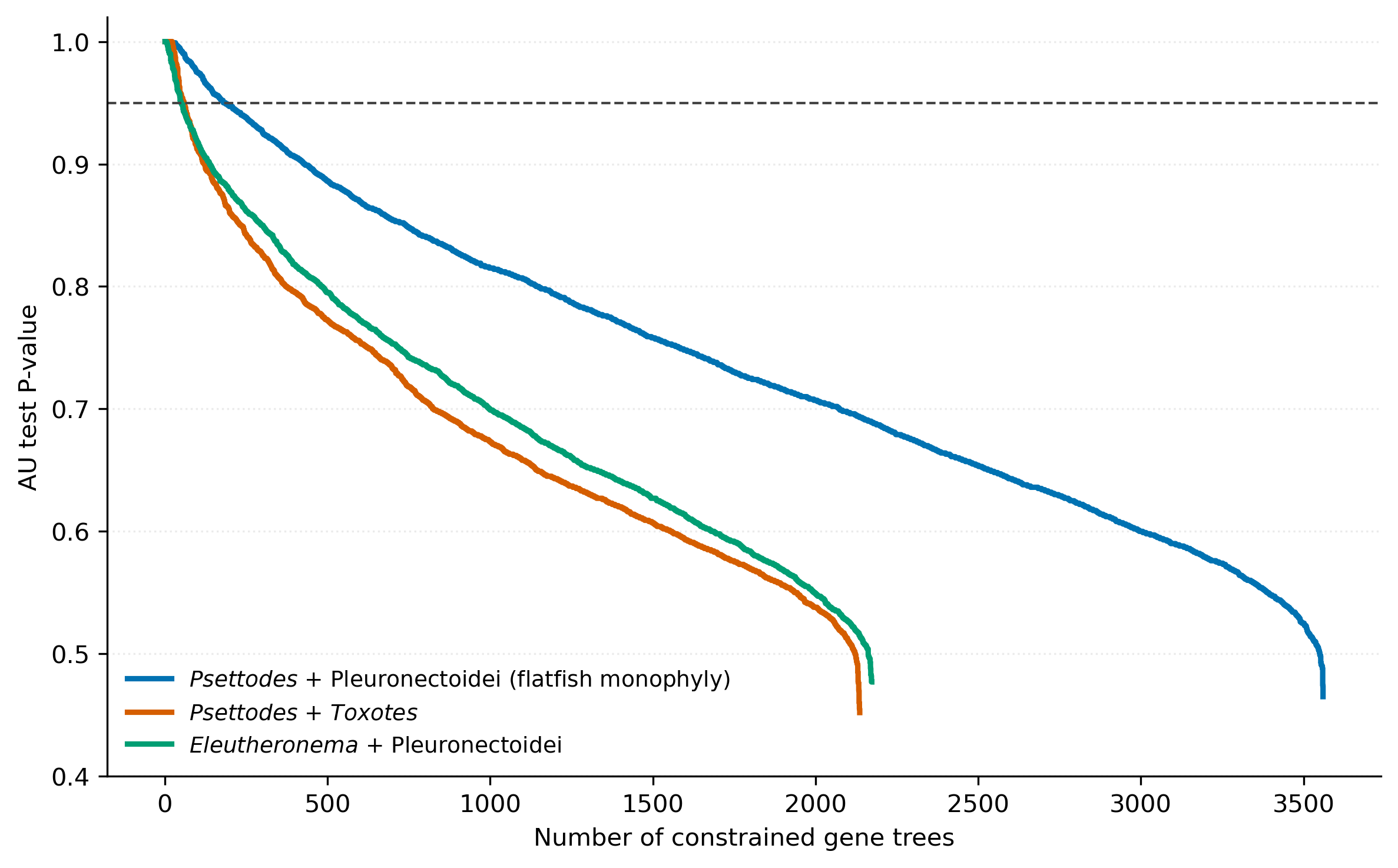

**Supplementary Figure S12.** AU-ranking curves across three constrained gene-tree topologies used in the per-gene approximately unbiased tests. Rank curves for the 7,869 genes jointly comparable across the three constrained topologies. The monophyly-compatible constraint is the top-ranked option for the largest share of loci, but most genes remain weakly discriminative among the alternatives.

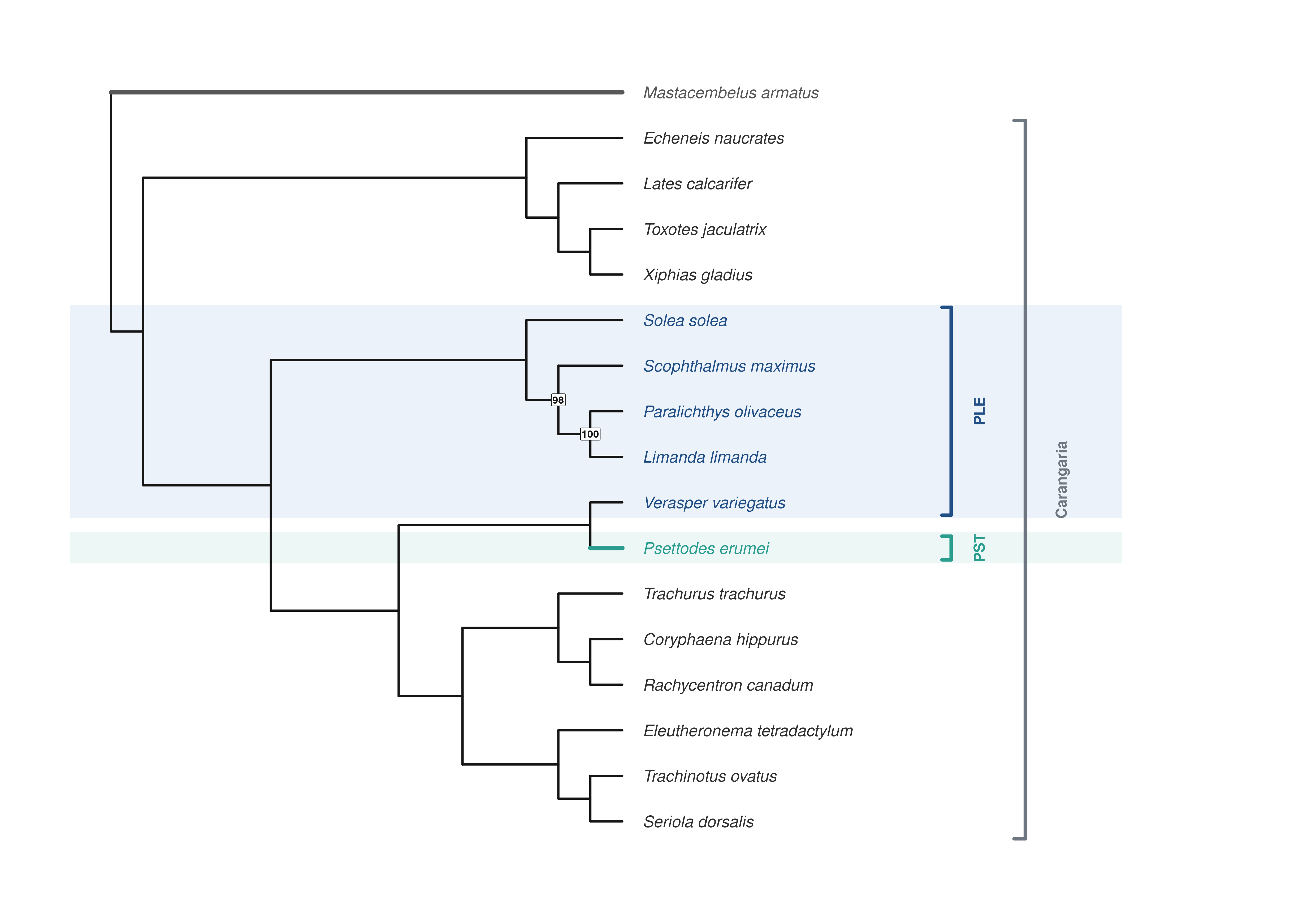

**Supplementary Figure S13.** RefSeq/New annotation-provenance microsynteny tree. Exploratory microsynteny phylogeny inferred with syntenet from a mixed annotation set in which species with available NCBI gene models were represented by RefSeq annotations and the remaining taxa by our newly generated annotations. Rather than recovering known phylogenetic relationships, species group by annotation source, indicating that heterogeneity in gene boundaries, exon structures, and isoform definitions—not biological signal—drives the inferred topology.

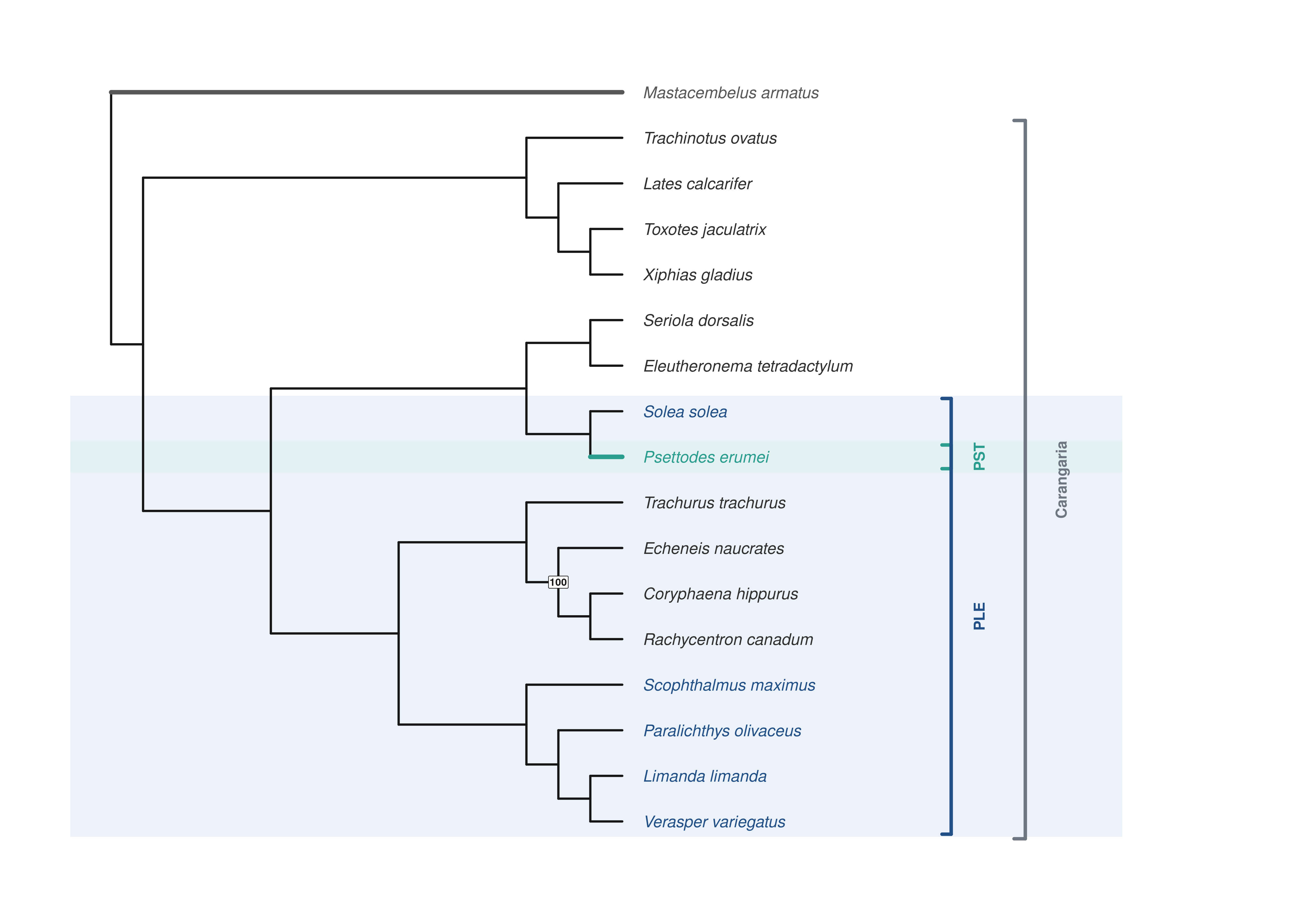

**Supplementary Figure S14.** BRAKER/GALBA annotation-provenance microsynteny tree. Exploratory microsynteny phylogeny inferred with syntenet from a second mixed-annotation regime in which some species were annotated only with BRAKER and others only with GALBA. As in Supplementary Figure S13, taxa cluster by annotation pipeline rather than by phylogenetic affinity, confirming that annotation provenance distorts protein-based synteny inference.

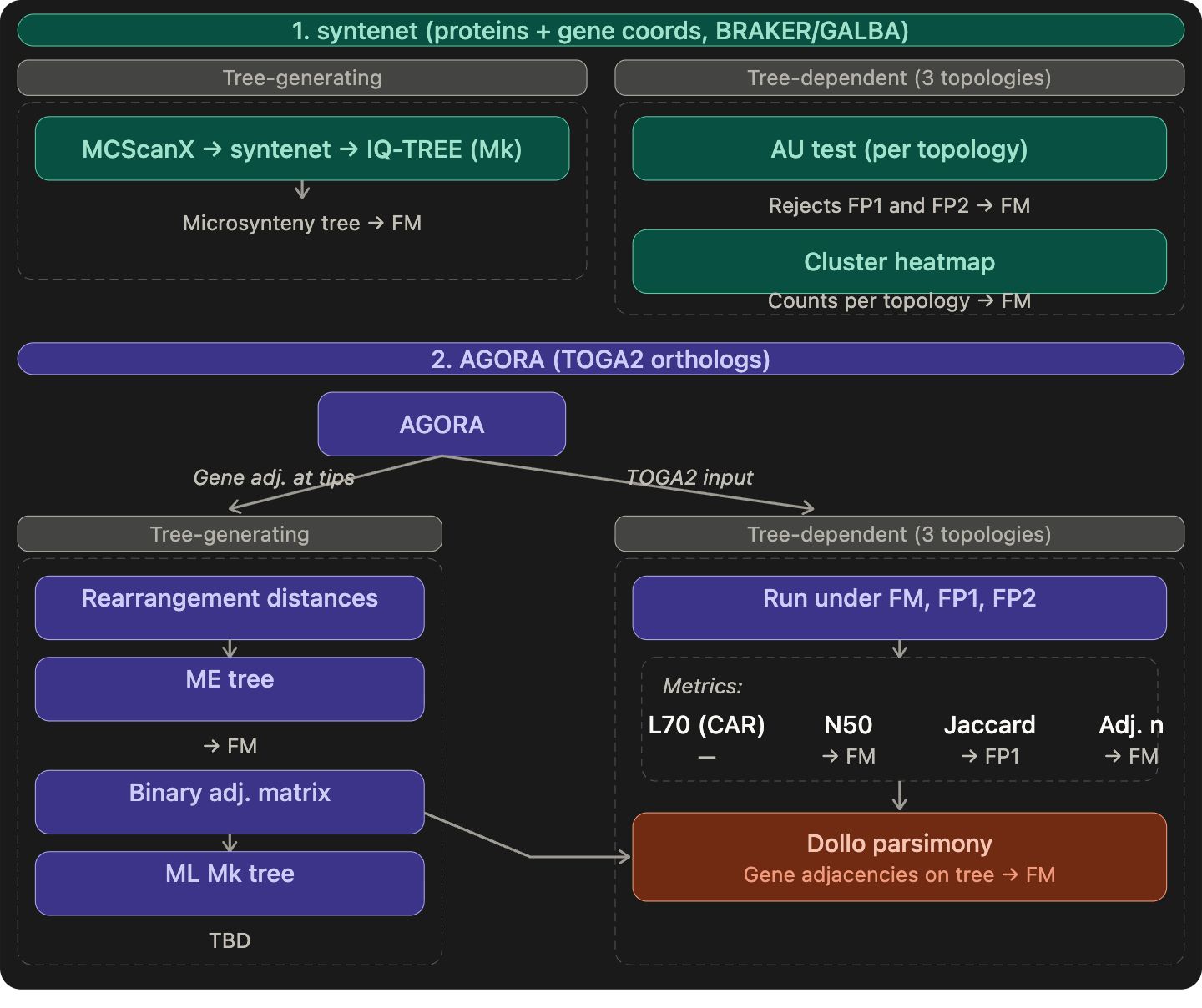

**Supplementary Figure S15.** Microsynteny analysis workflow summary. Flowchart summarizing the microsynteny analytical workflow, from harmonized annotation preparation and ortholog formatting through syntenet clustering, AGORA-based gene-order analyses, and downstream topology-testing summaries.

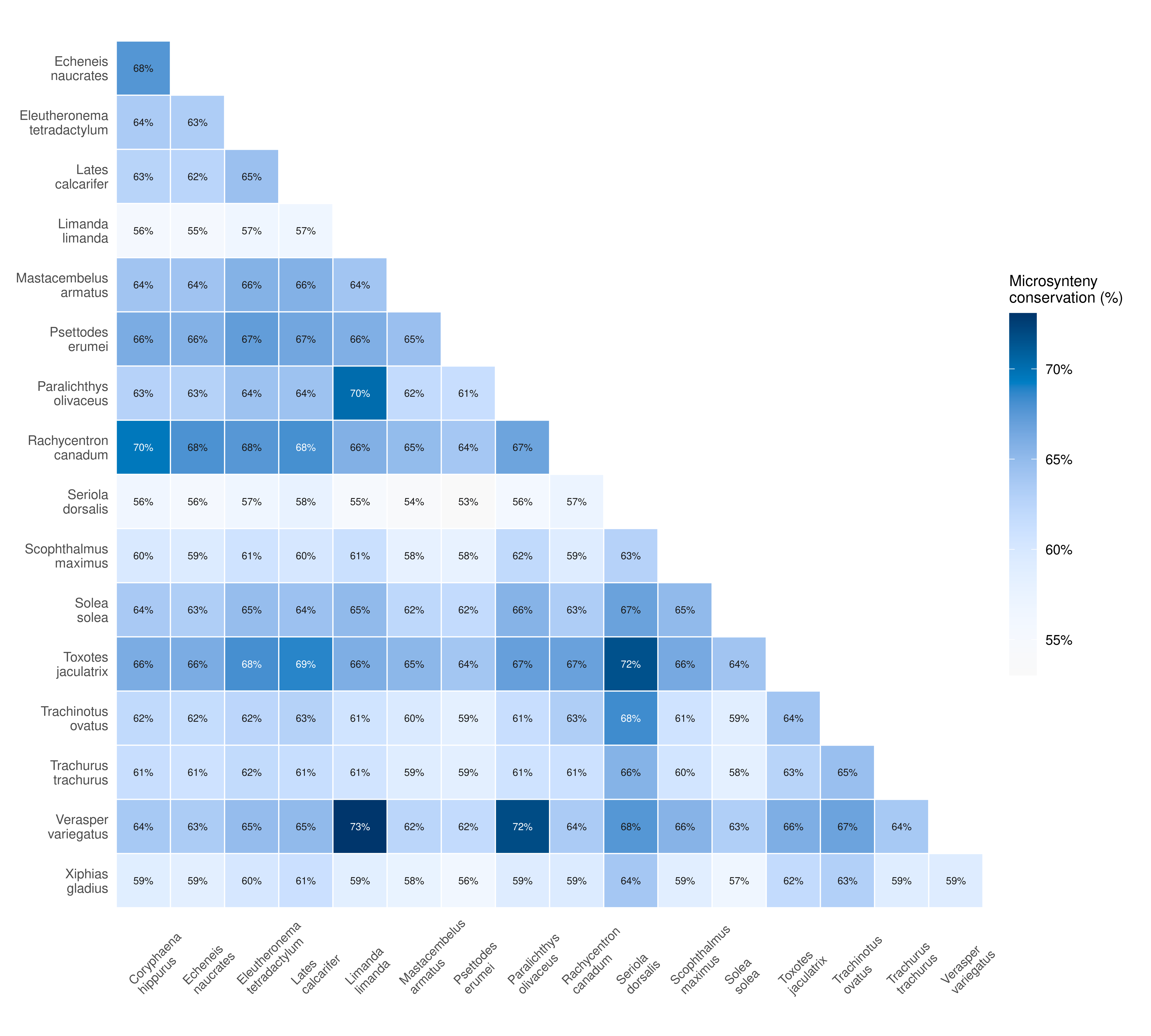

**Supplementary Figure S16.** Lower-triangular heatmap of pairwise microsynteny-conservation percentages among the 17 sampled genomes. Pairwise microsynteny-conservation matrix across the 17 sampled genomes. The strongest pairwise conservation occurs between *Verasper variegatus* and *Limanda limanda*, whereas one of the weakest values occurs between *Seriola dorsalis* and *Psettodes erumei*.

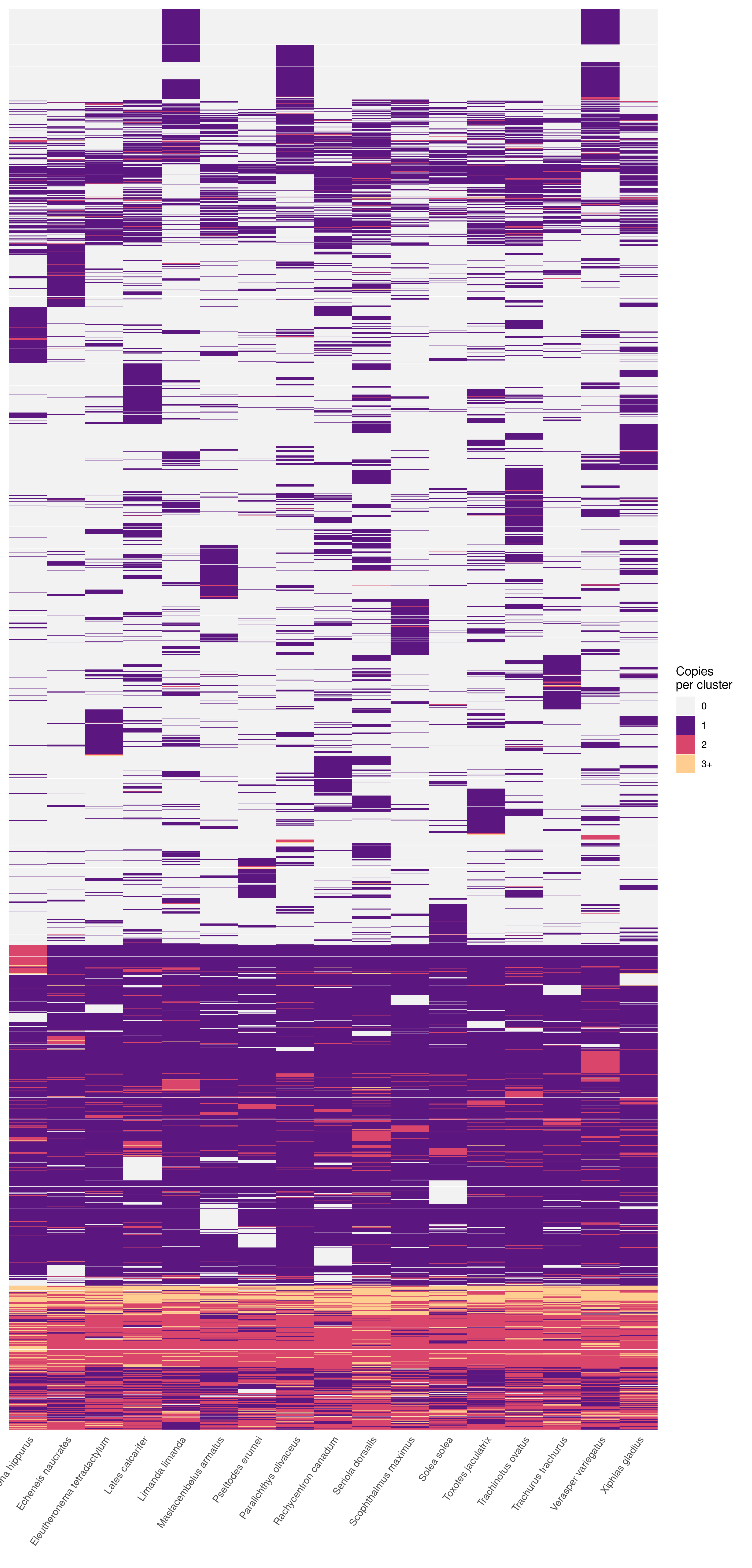

**Supplementary Figure S17.** Variant-only clustered microsynteny-profile heatmap summarizing synteny-cluster copy-number variation across the 17 sampled genomes. Variant-only view of the cluster-by-species microsynteny profile matrix after occupancy filtering and copy-number capping for visualization.

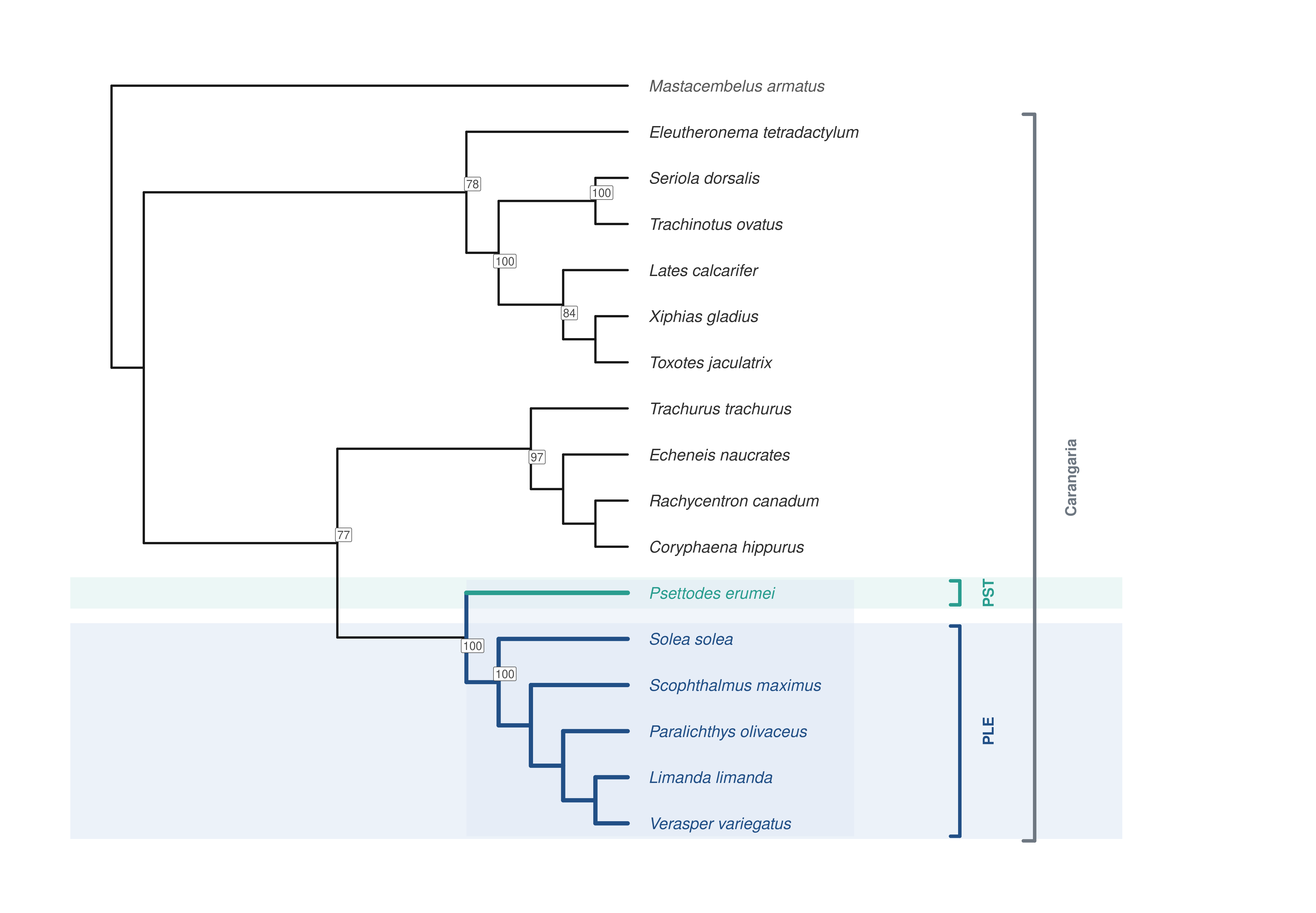

**Supplementary Figure S18.** Full-species microsynteny tree inferred from MCScanX collinearity with syntenet. Microsynteny-based tree for all 17 sampled species inferred with syntenet from MCScanX collinearity calls.

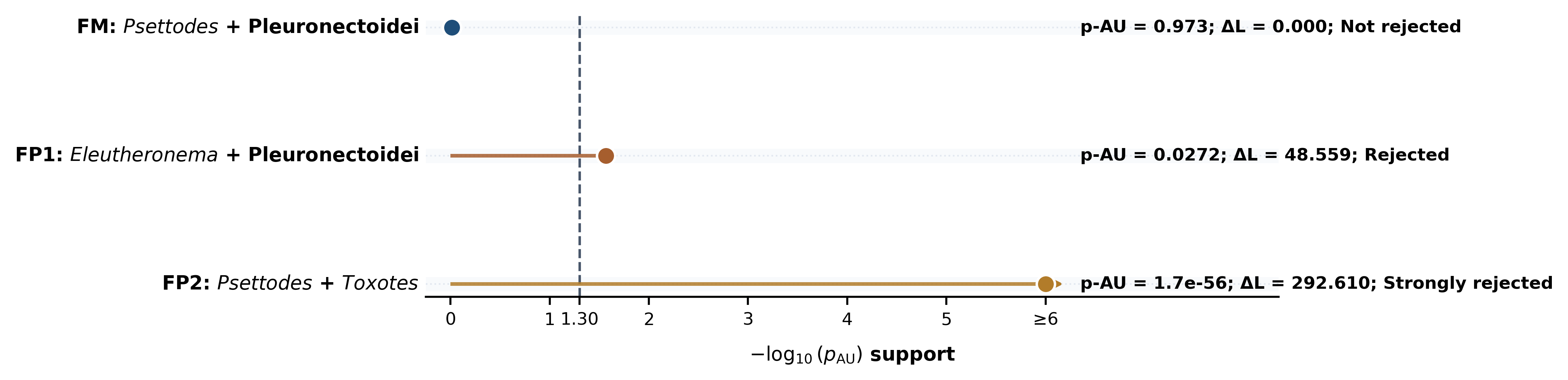

**Supplementary Figure S19.** Whole-matrix microsynteny AU-test summary across the three constrained topology hypotheses. Point summary of the three constrained microsynteny AU tests. FM is not rejected, FP1 is rejected, and FP2 is strongly rejected.

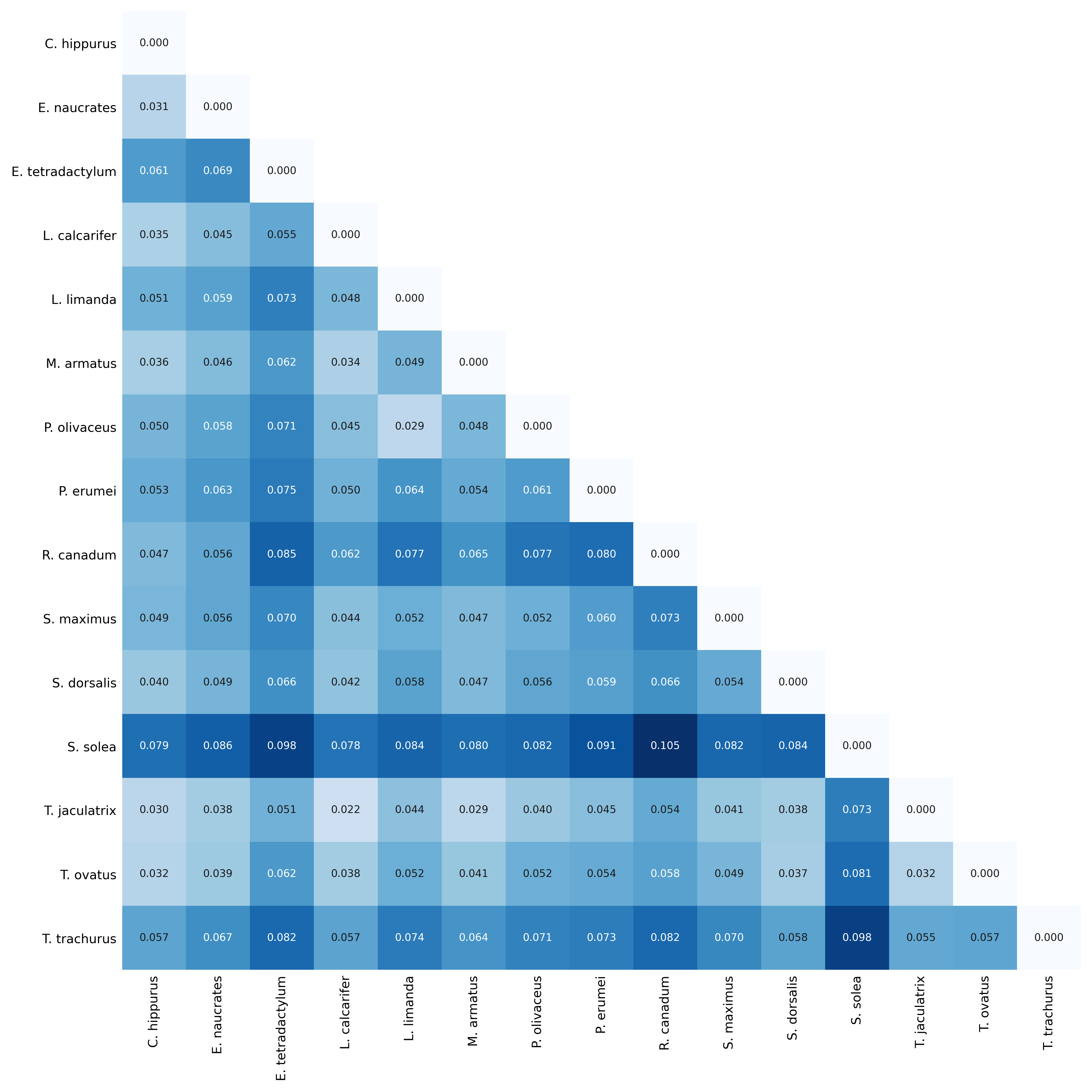

**Supplementary Figure S20.** Annotated heatmap of normalized pairwise adjacency distances for the primary strict full-species microsynteny dataset. Normalized adjacency-distance matrix for the strict 5,103-family full-species AGORA dataset.

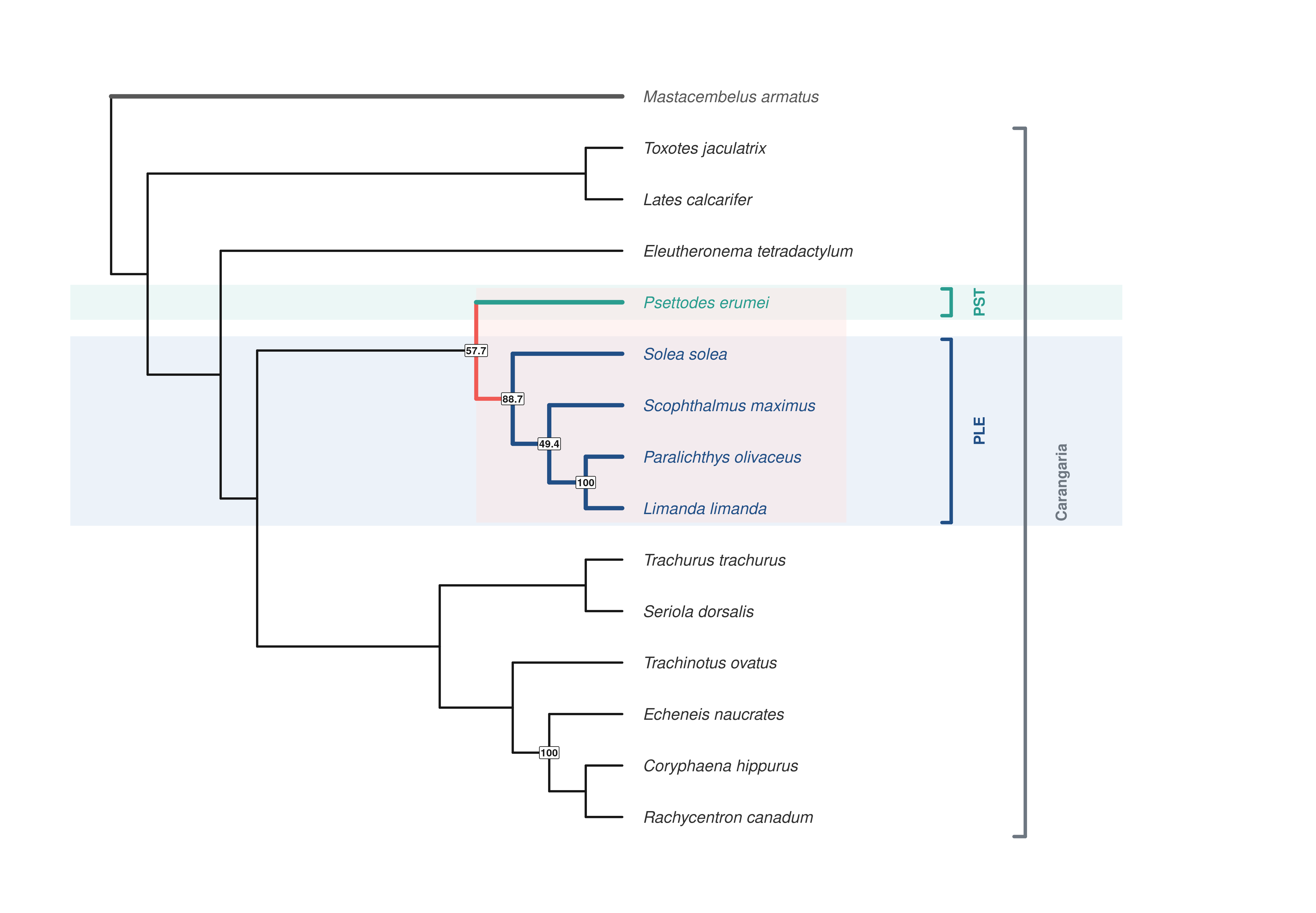

**Supplementary Figure S21.** Complete Minimum Evolution tree from the primary strict full-species microsynteny distance matrix. Complete Minimum Evolution tree inferred from the strict full-species adjacency-distance matrix.

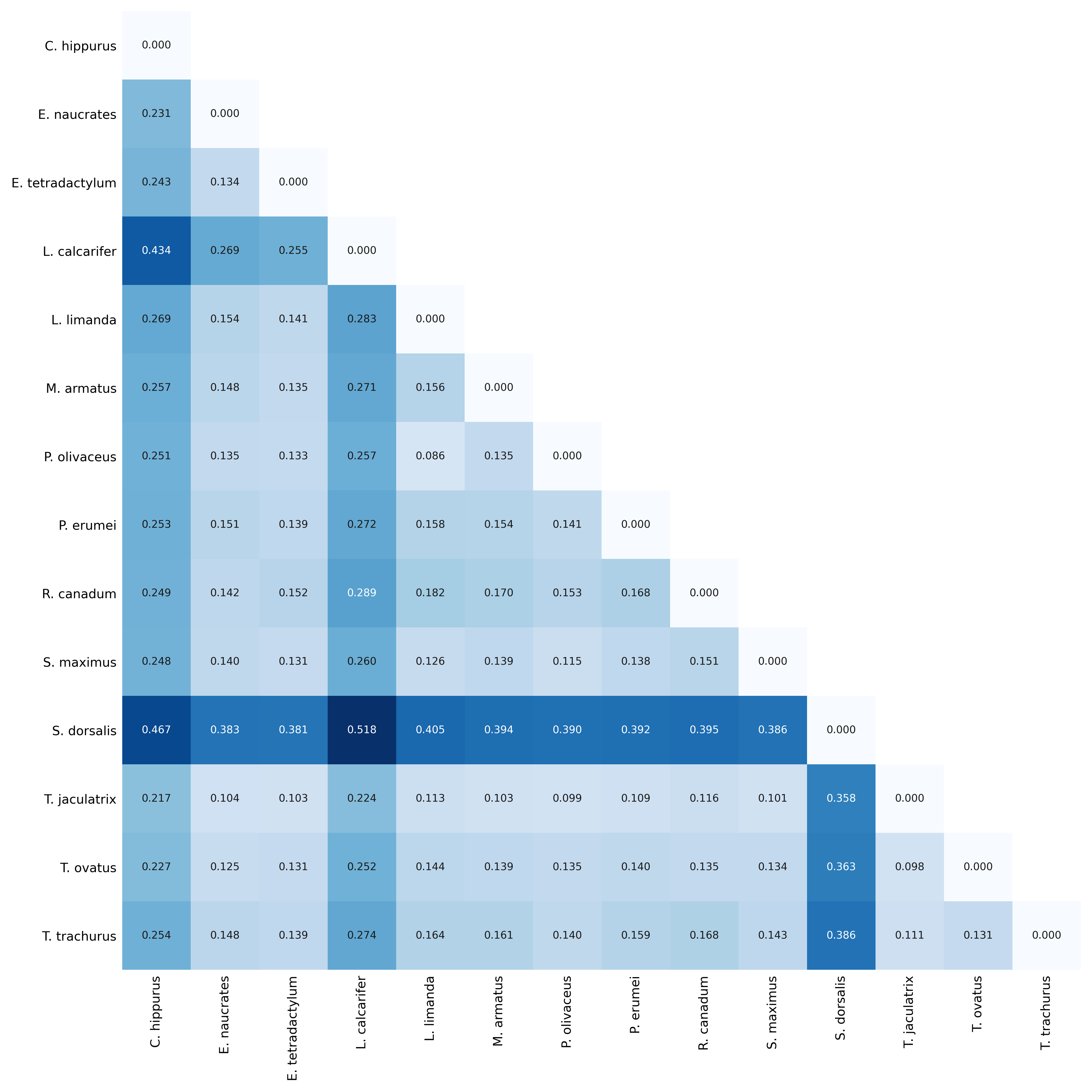

**Supplementary Figure S22.** Annotated heatmap of normalized pairwise adjacency distances for the no-Solea microsynteny sensitivity dataset. Normalized adjacency-distance matrix after removal of *Solea solea* from the strict AGORA sensitivity dataset.

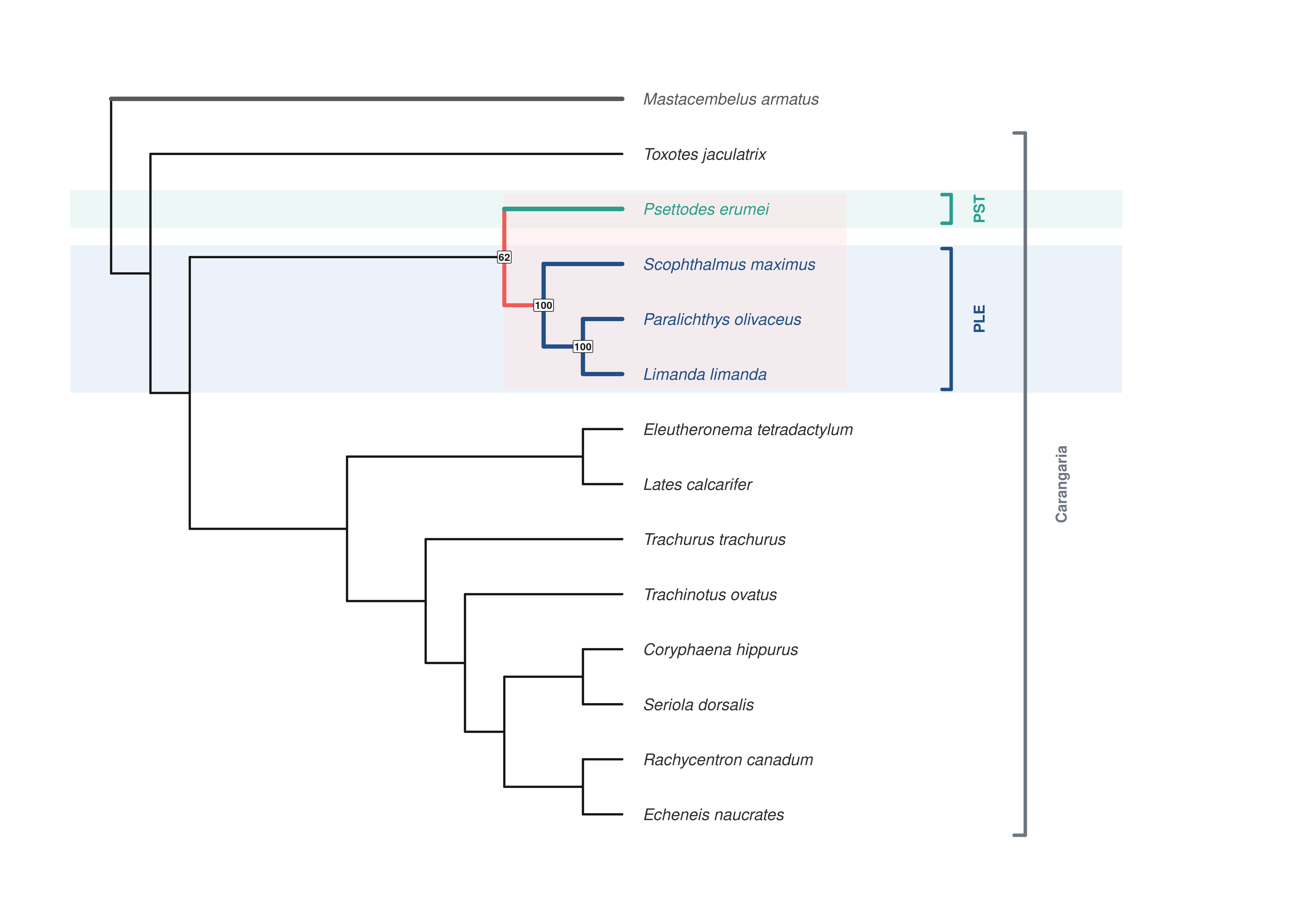

**Supplementary Figure S23.** Complete Minimum Evolution tree from the no-Solea microsynteny distance matrix. Complete Minimum Evolution tree inferred from the no-Solea sensitivity matrix.

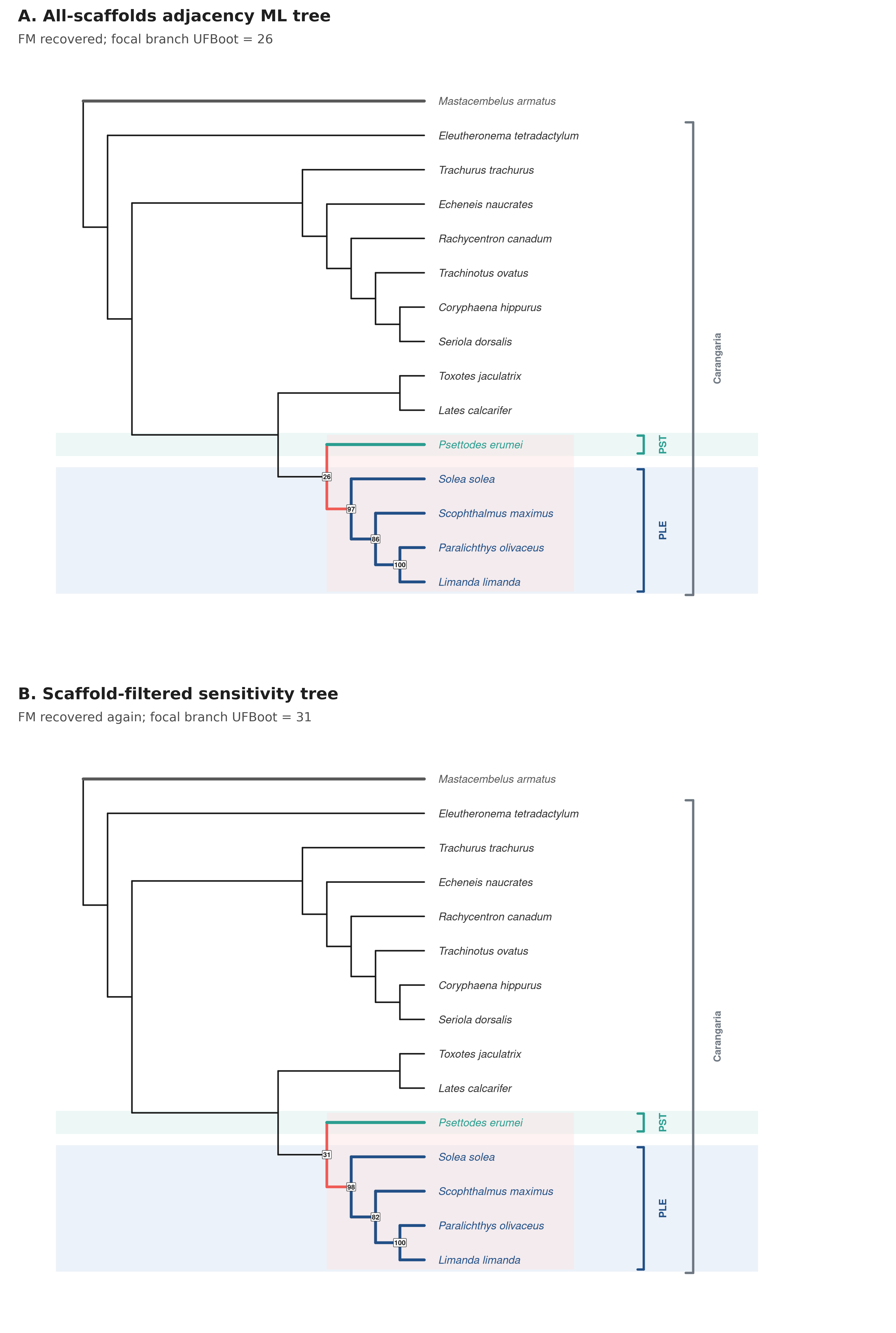

**Supplementary Figure S24.** Binary adjacency maximum-likelihood trees from the all-scaffolds and scaffold-filtered datasets. Two-panel summary of the IQ-TREE binary gene-adjacency analyses described in Supplementary Table S14. Panel A shows the all-scaffolds analysis from the 0.99 ortholog set; panel B shows the sensitivity analysis restricted to orthogroups on scaffolds with at least 20 mapped markers. In both trees, *Psettodes erumei* is recovered as sister to the sampled Pleuronectoidei; node labels are SH-aLRT/UFBoot percentages, with focal FM support of 71.2/26 in panel A and 70.3/31 in panel B.

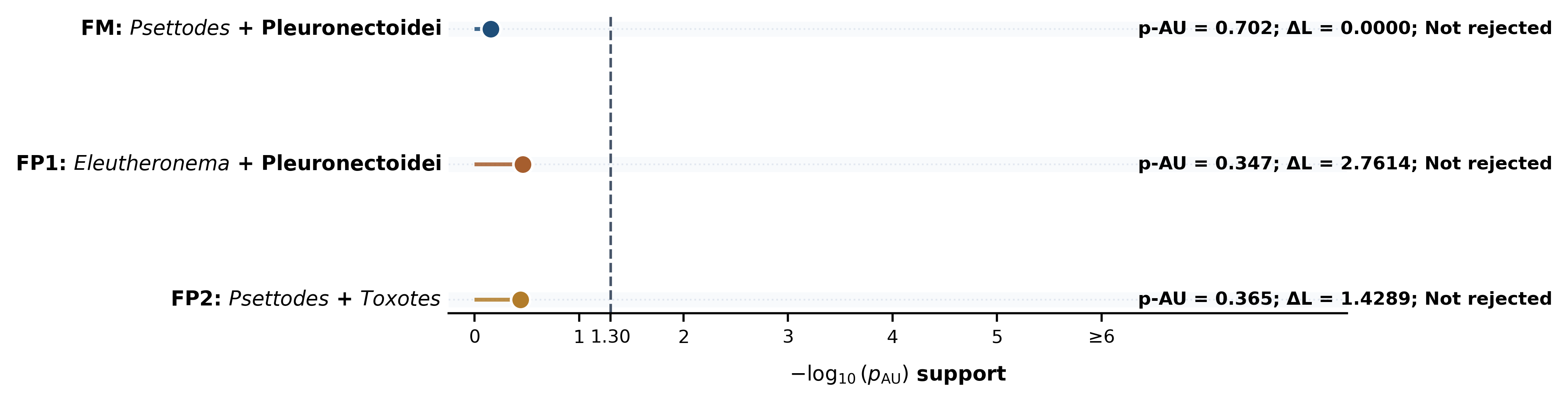

**Supplementary Figure S25.** Binary adjacency AU-test summary across the three constrained topology hypotheses. Point summary of the constrained IQ-TREE AU tests on the all-scaffolds binary adjacency matrix summarized in Supplementary Table S14. FM had the highest likelihood (deltaL = 0; p-AU = 0.702), while FP1 (deltaL = 2.7614; p-AU = 0.347) and FP2 (deltaL = 1.4289; p-AU = 0.365) also remained in the AU confidence set. The dashed vertical line marks the conventional p-AU = 0.05 rejection threshold.

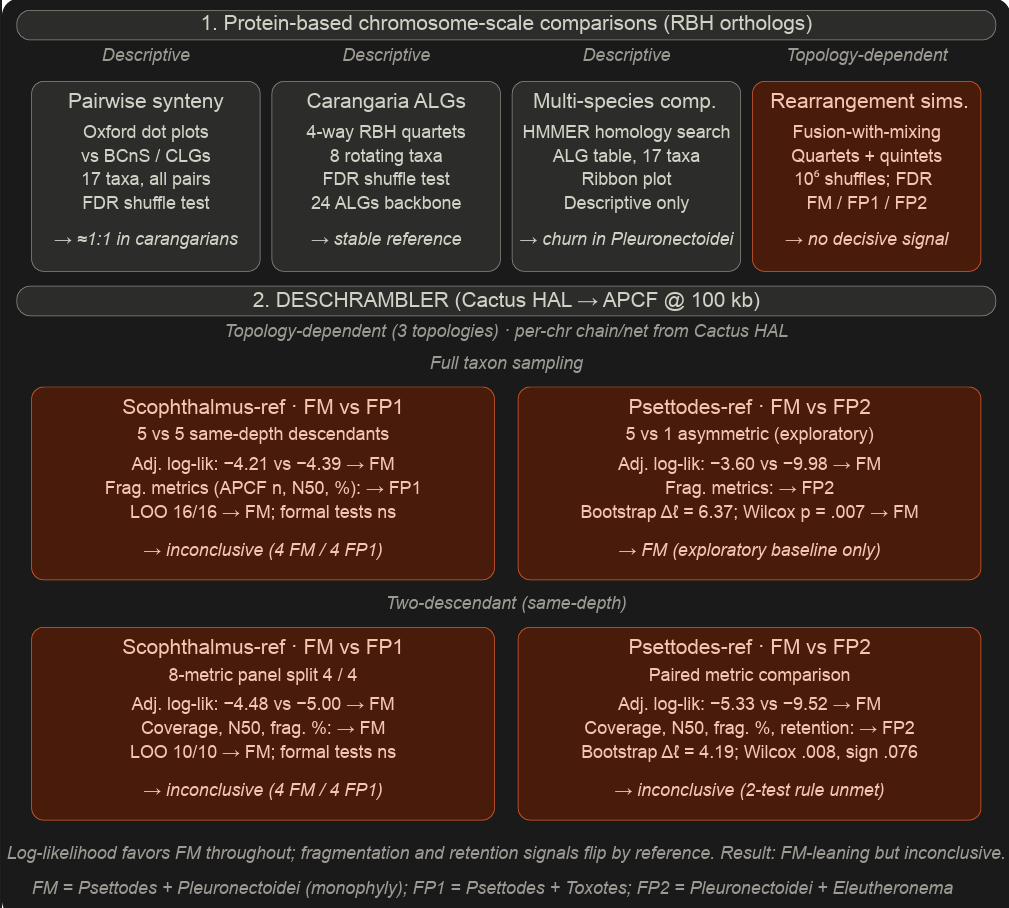

**Supplementary Figure S26.** Macrosynteny analysis workflow summary. Flowchart summarizing the macrosynteny analytical workflow, from pairwise chromosome-scale reciprocal-best-hit comparisons and ancestral linkage-group definition through rearrangement simulations and DESCHRAMBLER-based ancestral chromosome reconstruction.

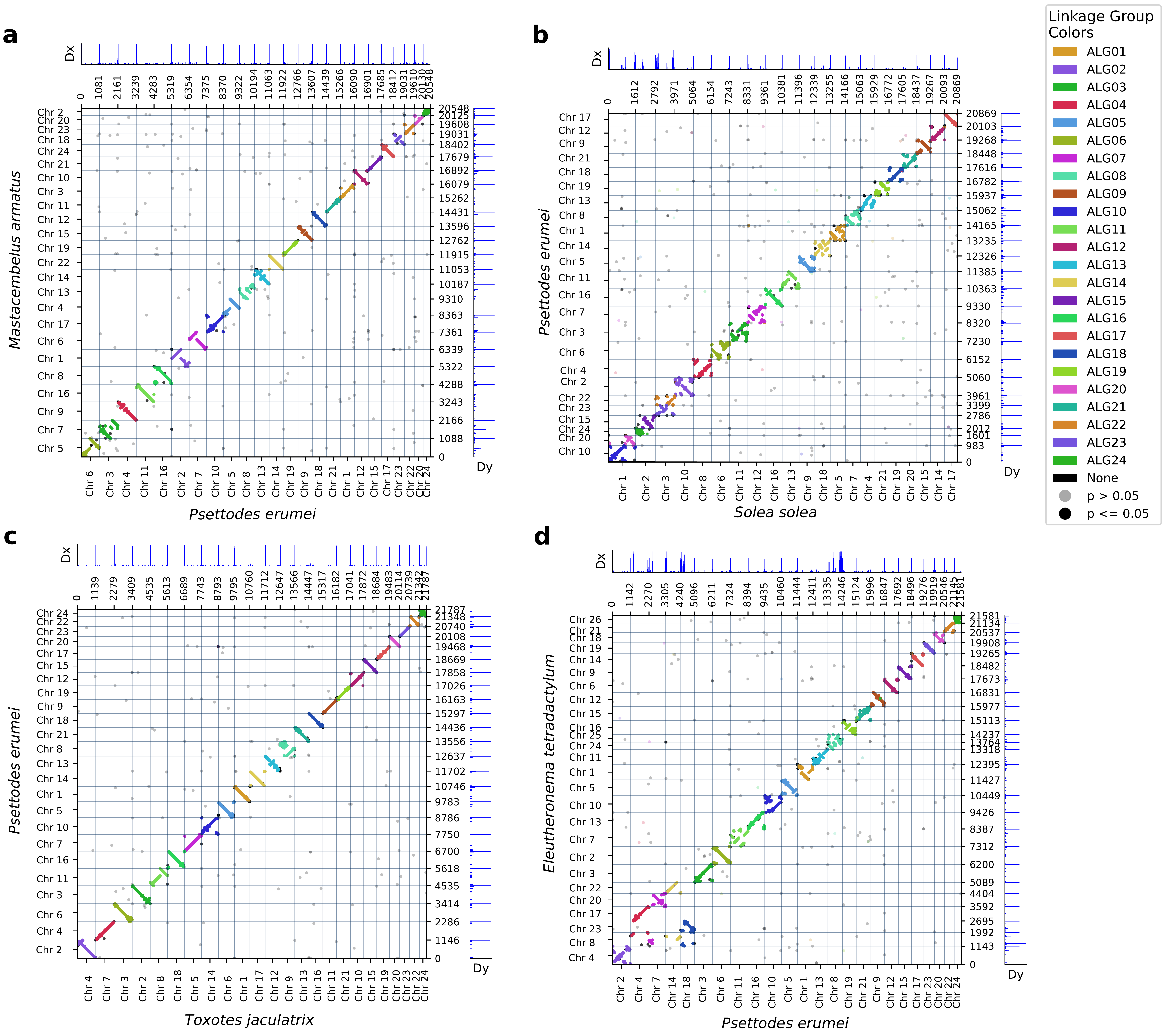

**Supplementary Figure S27.** Macrosynteny chromosome-comparison oxford dot plots for *Psettodes erumei*. Representative Oxford dot plots illustrating chromosome-scale orthologous relationships between *Psettodes erumei* and four key comparative taxa. Panels show macrosynteny conservation and structural rearrangements relative to (a) the outgroup *Mastacembelus armatus*; (b) *Solea solea*; (c) *Toxotes jaculatrix*; and (d) *Eleutheronema tetradactylum*. *Psettodes erumei* retains predominantly one-to-one chromosome correspondences with non-flatfish carangarians, consistent with retention of the ancestral carangarian karyotype. The extensive fusion, fission, and segment mixing seen against the pleuronectoid comparator (Solea solea), together with the lineage-specific chromosome fission distinguishing the polynemid Eleutheronema tetradactylum, are therefore independently derived features.

**Panel A.** FM topology (*Psettodes erumei* + Pleuronectoidei).

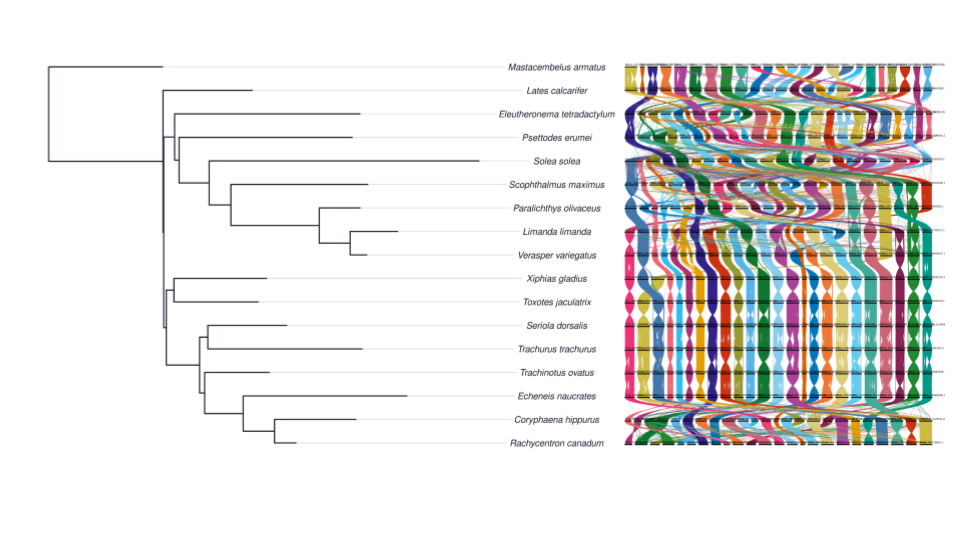

**Panel B.** FP1 topology (*Eleutheronema tetradactylum* + Pleuronectoidei).

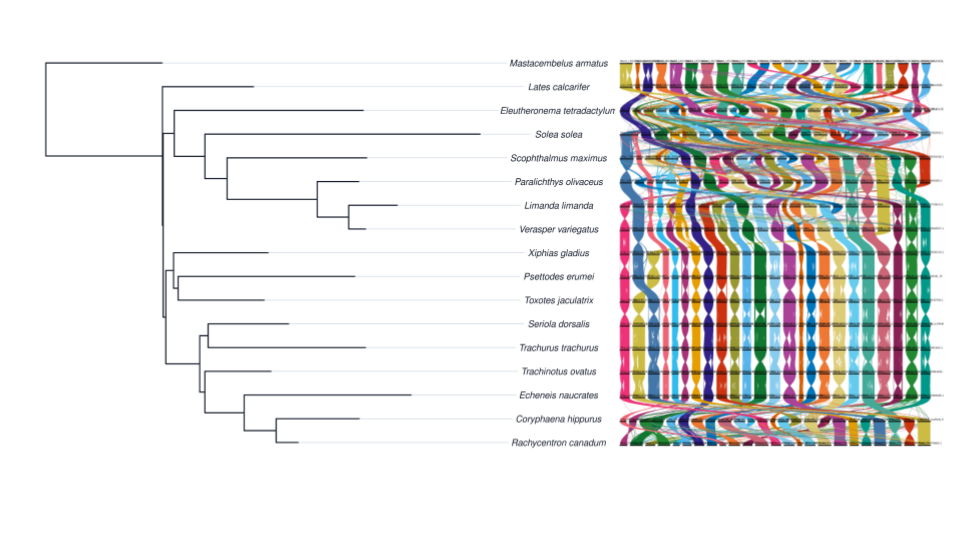

**Panel C.** FP2 topology (*Psettodes erumei* + *Toxotes jaculatrix*).

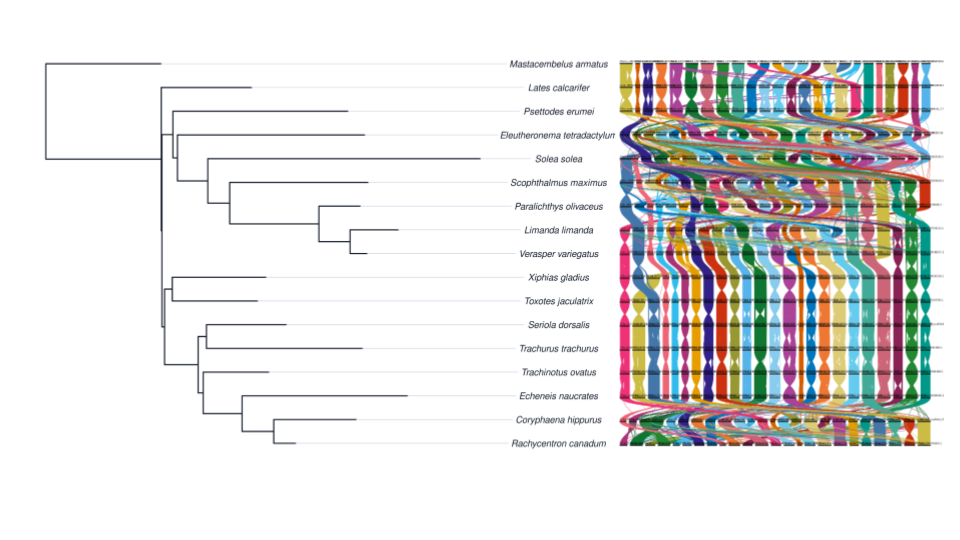

**Supplementary Figure S28.** Macrosynteny ribbon plots under the three alternative topology hypotheses. Composite macrosynteny ribbon panels tracing ALG01-ALG24 across the sampled genomes under each of the three alternative focal topologies. In each panel, the tree at left shows the corresponding constrained relationship, and the ribbon plot at right shows the same multispecies ALG correspondences in the matching taxon order to facilitate direct comparison of large-scale chromosome-structure patterns among the FM, FP1, and FP2 hypotheses.

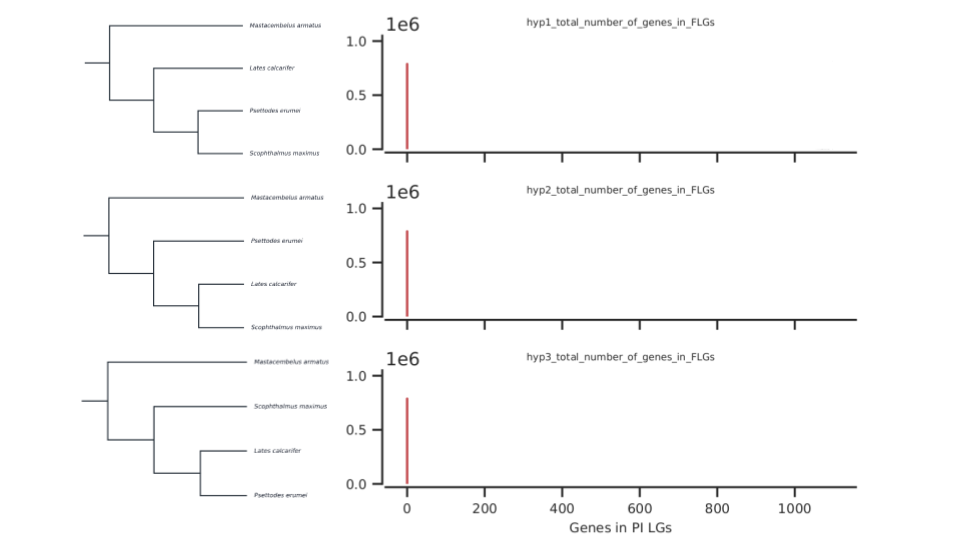

**Supplementary Figure S29.** Quartet genome rearrangement simulation summary for FM-compatible taxon sets. Quartet-based fusion-with-mixing simulation panel for the FM-compatible taxon configurations. Trees at left show the alternative quartet hypotheses evaluated against the same outgroup-rooted taxon set, and the summary panels at right show the corresponding topology-support scoring output under genome-label randomization.

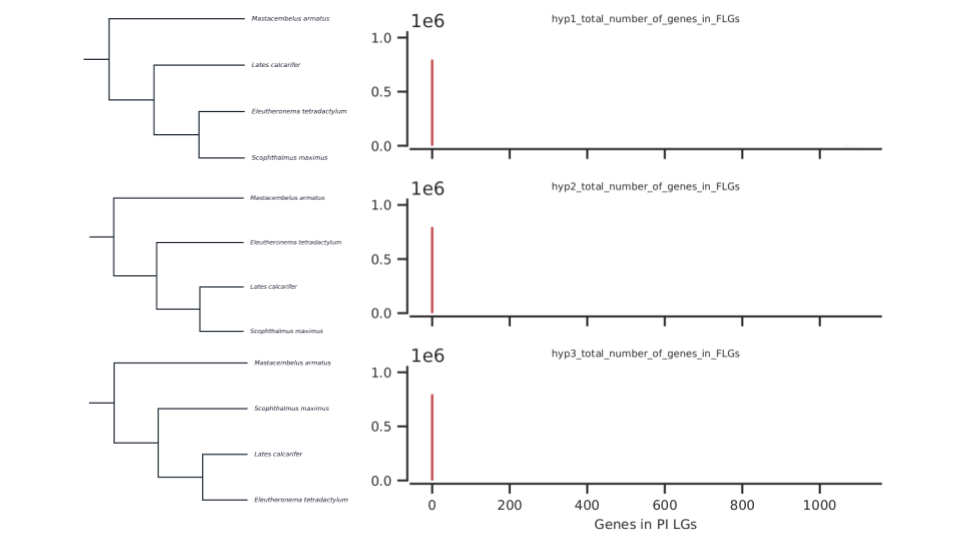

**Supplementary Figure S30.** Quartet genome rearrangement simulation summary for FP1-compatible taxon sets. Quartet-based fusion-with-mixing simulation panel for the FP1-compatible taxon configurations, evaluated against the same three alternative quartet hypotheses under the randomization framework described in the methods.

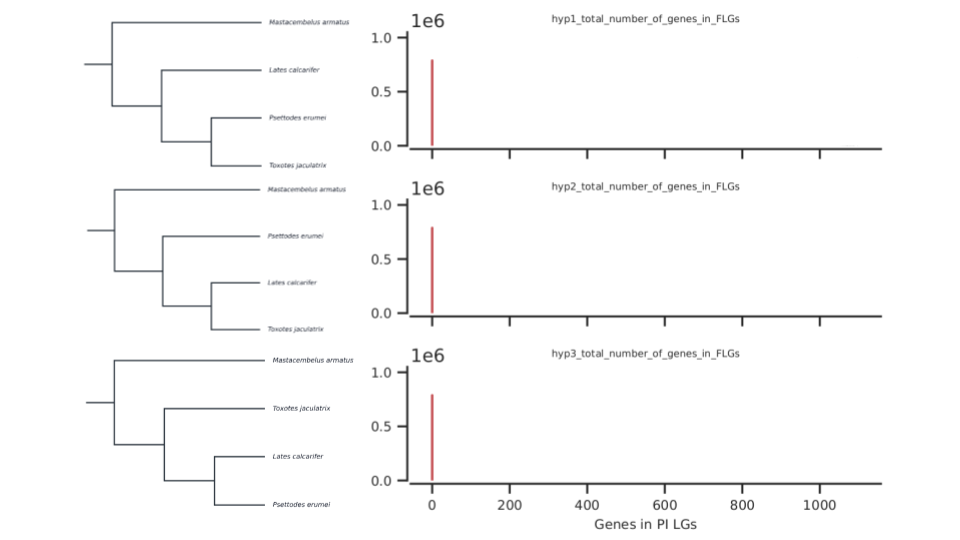

**Supplementary Figure S31.** Quartet genome rearrangement simulation summary for FP2-compatible taxon sets. Quartet-based fusion-with-mixing simulation panel for the FP2-compatible taxon configurations. Together with Supplementary Figures S28-S32, this figure shows that no quartet or quintet design provided decisive support for a single focal topology.

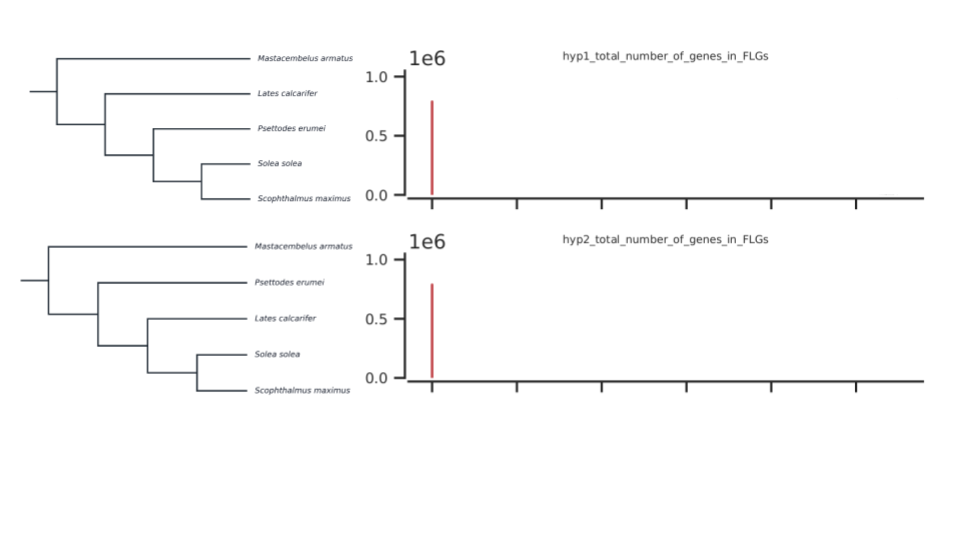

**Supplementary Figure S32.** Quintet genome rearrangement simulation summary for FM-compatible taxon sets. Quintet-based fusion-with-mixing simulation panel for the FM-compatible taxon configuration with two fixed pleuronectoid taxa. The two trees at left show the only alternative quintet hypotheses evaluated for this shared outgroup-rooted taxon set, and the summary panels at right show the corresponding topology-support scoring output under genome-label randomization.

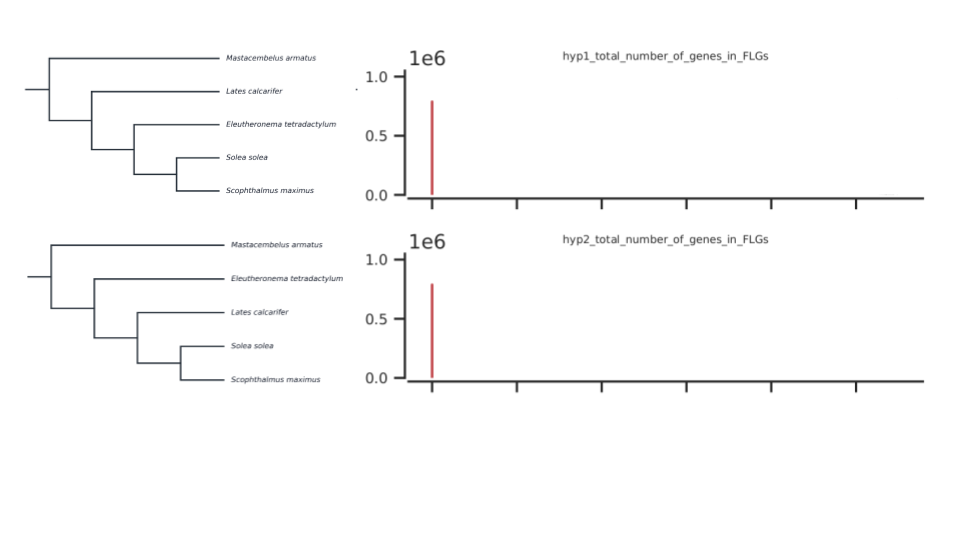

**Supplementary Figure S33.** Quintet genome rearrangement simulation summary for FP2-compatible taxon sets. Quintet-based fusion-with-mixing simulation panel for the FP2-compatible taxon configuration with two fixed pleuronectoid taxa. As in Supplementary Figure S32, only the two alternative quintet hypotheses compatible with the fixed pleuronectoid pair were tested, and the right-hand panels summarize their observed topology-support scores under genome-label randomization.

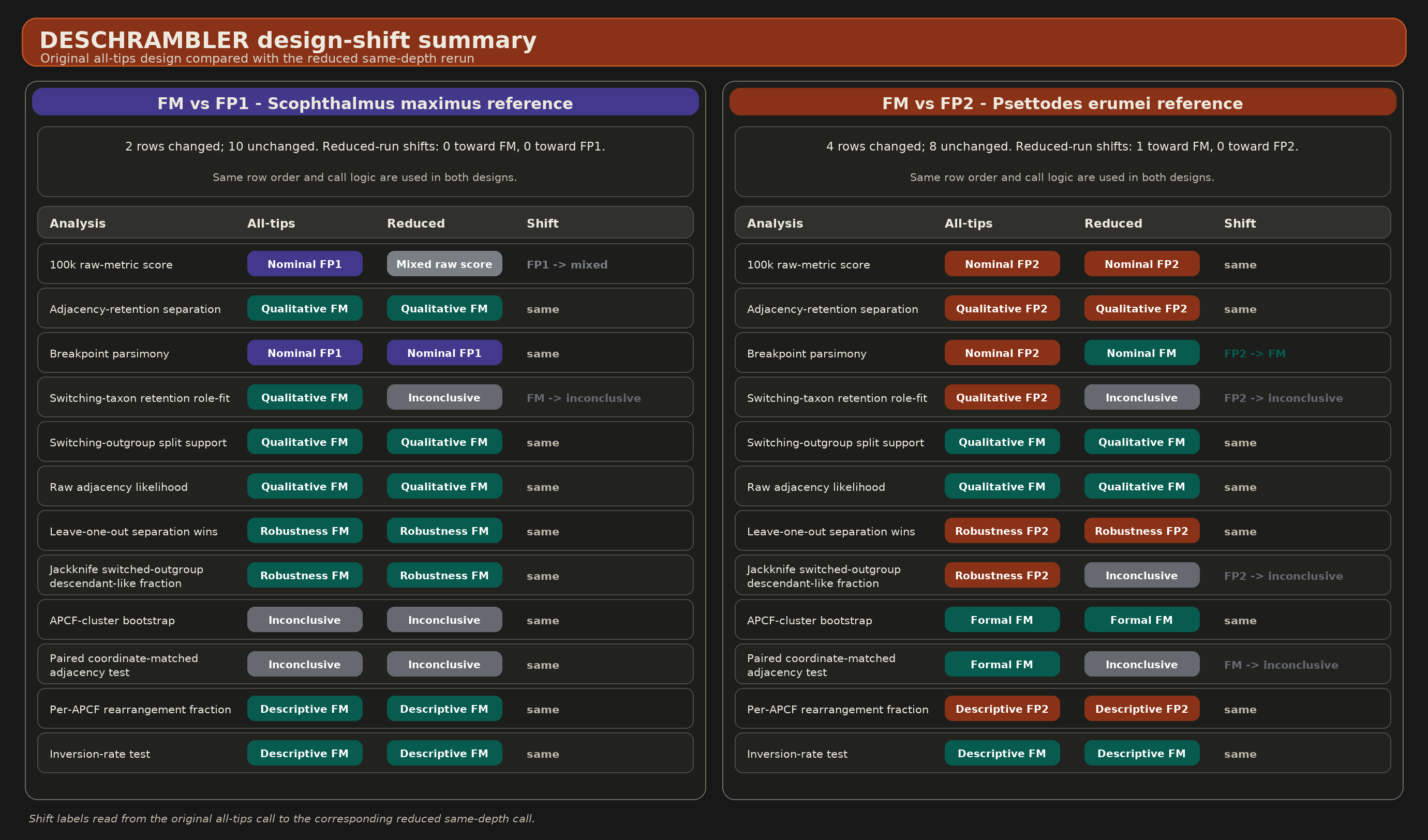

**Supplementary Figure S34.** DESCHRAMBLER design-shift summary from the all-tips design to the reduced same-depth design. Comparison of row-level DESCHRAMBLER call shifts from the original all-tips design to the reduced same-depth rerun across the FM vs FP1 and FM vs FP2 shared-reference comparisons.

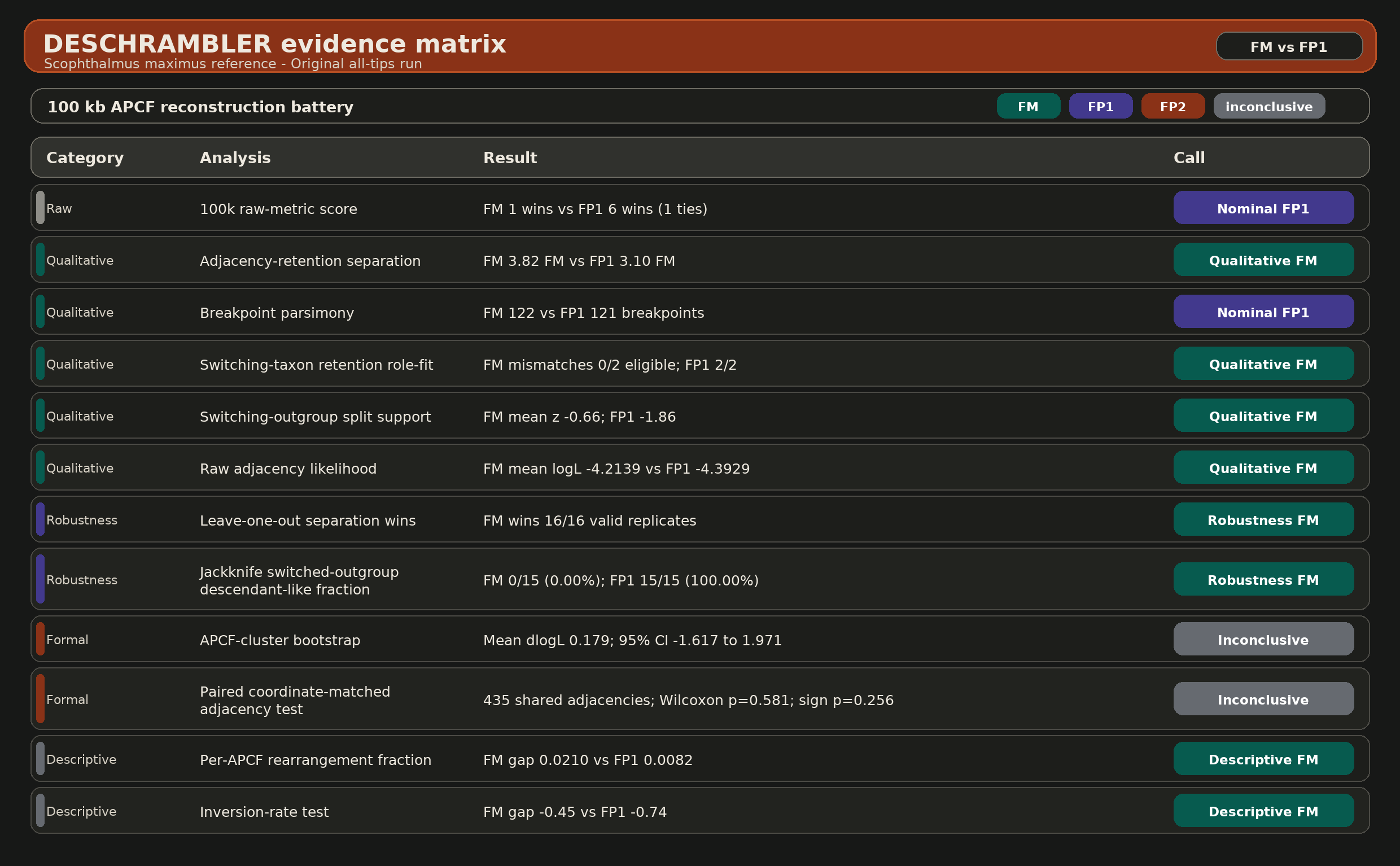

**Supplementary Figure S35.** DESCHRAMBLER evidence matrix for the all-tips FM vs FP1 comparison. Evidence matrix for the original all-tips DESCHRAMBLER comparison of FM versus FP1 under the *Scophthalmus maximus* reference. Each row summarizes one component of the shared 100-kb comparison battery and indicates whether the corresponding signal nominally favors FM, favors FP1, or remains inconclusive.

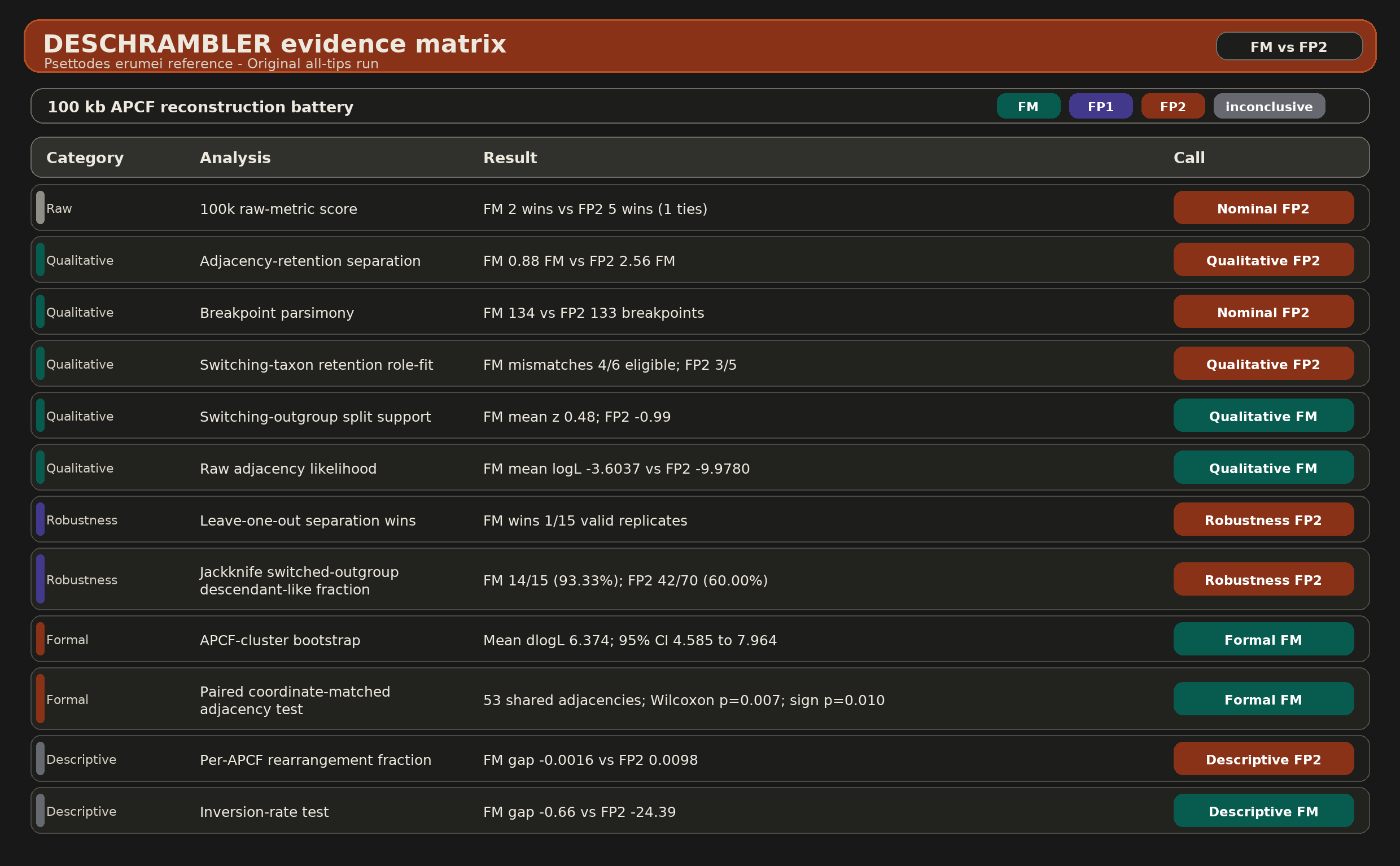

**Supplementary Figure S36.** DESCHRAMBLER evidence matrix for the all-tips FM vs FP2 comparison. Evidence matrix for the original all-tips DESCHRAMBLER comparison of FM versus FP2 under the *Psettodes erumei* reference. The panel summarizes the same 100-kb analysis battery used in Supplementary Figure S35, allowing direct comparison of which signals favor FM, favor FP2, or remain inconclusive in the mixed-depth design.

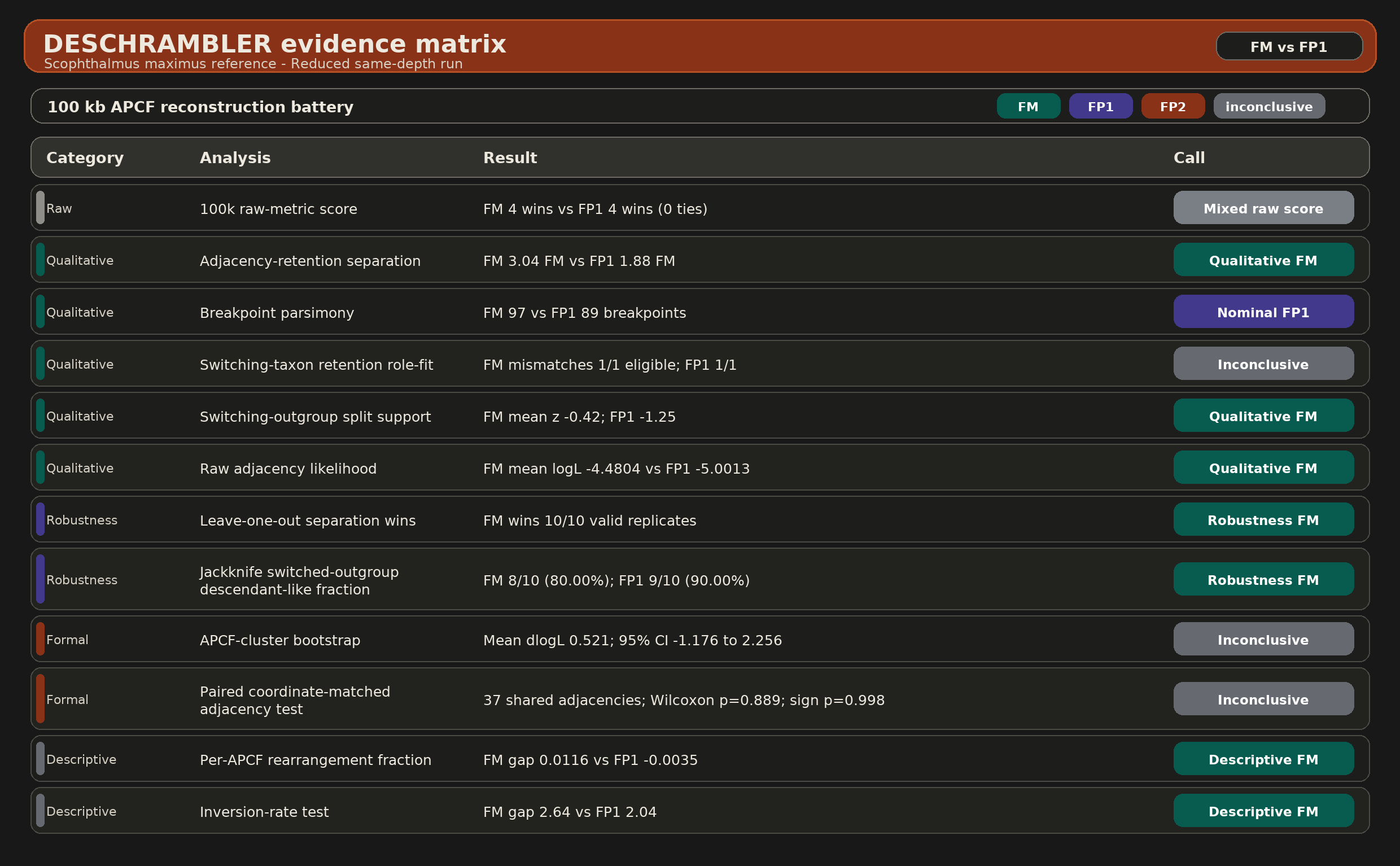

**Supplementary Figure S37.** DESCHRAMBLER evidence matrix for the reduced same-depth FM vs FP1 comparison. Evidence matrix for the reduced same-depth DESCHRAMBLER rerun comparing FM and FP1 under the *Scophthalmus maximus* reference. By holding ingroup depth constant across the shared-reference runs, this panel isolates how the same 100-kb evidence battery behaves after removing the original mixed-depth asymmetry.

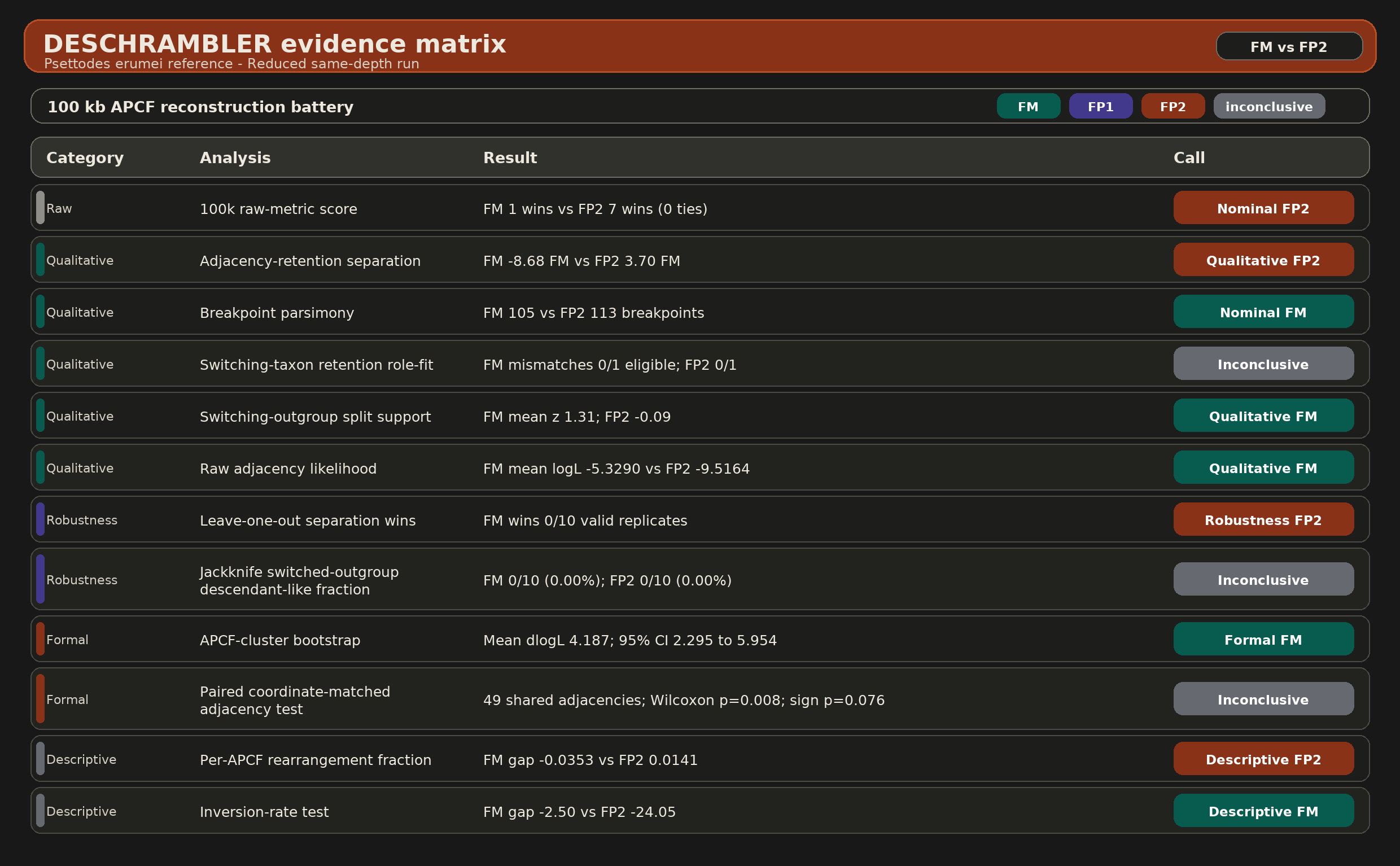

**Supplementary Figure S38.** DESCHRAMBLER evidence matrix for the reduced same-depth FM vs FP2 comparison. Evidence matrix for the reduced same-depth DESCHRAMBLER rerun comparing FM and FP2 under the *Psettodes erumei* reference. Together with Supplementary Figures S34-S36, this panel provides the pairwise evidence tables for both shared-reference comparisons in the original all-tips and balanced same-depth designs.
